# High-light adaptation in *Synechocystis* by accumulating NDH proteins and depleting specific phycobilisome linker proteins

**DOI:** 10.1101/2025.07.21.665844

**Authors:** Weiyang Chen, Eslam M. Abdel-Salam, Marcel Dann, Caroline Ott, Serena Schwenkert, Dario Leister

## Abstract

Photosynthetic organisms have evolved mechanisms to manage excess light, crucial for maximizing photosynthetic efficiency. High-light (HL) tolerant *Synechocystis* sp. PCC6803 strains were developed through laboratory evolution, with tolerance attributed to specific point mutations. Key mutations affected the NDH-1L complex F1-subunit (NdhF1_F124L_) and translation elongation factor G2 (EF-G2_R461C_). Reintroducing these mutations into laboratory strains conferred HL tolerance. Comparisons with knockout and overexpressor lines showed NdhF1_F124L_ and EF-G2_R461C_ result in gain of function. Transcriptomic and proteomic analysis unveiled a network of responses contributing to HL tolerance, including maintenance of phosphate metabolism and decreased antenna size by depleting a specific linker protein in EF-G2_R461C_ cells. Consequently, overexpression of Pho regulon genes increased HL tolerance. NdhF1_F124L_ enhances cyclic electron flow (CEF) by increasing NDH-1 complex subunit accumulation. Other HL-adapted strains demonstrated that increased CEF and decreased antenna size are recurring outcomes, achievable through various mutations. This study demonstrates how limited mutations can reconfigure cells for enhanced HL tolerance, offering insights for improving photosynthetic efficiency.

## Introduction

Improving photosynthesis is crucial for addressing future food, energy, and climate challenges (*1, 2*). One promising approach involves broadening the absorption spectrum through synthetic biology and reconfiguring photosynthetic light reactions (*3–5*). However, excessive solar energy input can cause photodamage (*6*), necessitating the development of new chassis with strong resilience to high light (HL).

Cyanobacteria serve as excellent models for manipulating photosynthesis (*7, 8*). Previous research using adaptive laboratory evolution (ALE) generated *Synechocystis* sp. PCC6803 (hereafter *Synechocystis*) strains capable of surviving extreme HL conditions (*9*). Two key mutations, phenylalanine (F) 124 to leucine (L) in the F1 subunit of the type 1 NAD(P)H dehydrogenase (NDH-1) complex (referred to as NdhF1_F124L_) and arginine (R) 461 to cysteine (C) in the translation elongation factor EF-G2 (hereafter referred to as EF-G2_R461C_), were found to increase HL tolerance in laboratory-type (LT) background (*9*).

NdhF exists in three isoforms, each incorporated into distinct NDH-1 complexes (*10, 11*). The NDH-1L complex, which includes NdhF1, is predominant under normal growth conditions and facilitates cyclic electron flow (CEF) around photosystem I (PSI) during short-term HL exposure and respiration (*11, 12*). In contrast, NdhF3 and NdhF4 are components of the NDH-1MS complex, which is involved in CO_2_ uptake (*11, 12*) and further enhances CEF under extended HL stress periods (*13, 14*). CEF plays a vital role in protecting cyanobacteria from reactive oxygen species (ROS) damage during environmental stress by preventing excess electron accumulation at the PSI acceptor side and generating additional ATP (*12*). Previous studies have demonstrated that HL exposure significantly induces NDH-1 subunit expression (*14–16*). Additionally, our prior research revealed that the NdhF1_F124L_ mutation enhances CEF activity, protecting cells against HL stress, shifting the ratio of CEF to CO_2_ uptake activities in favor of CEF, and boosting respiration (*9*).

The translation elongation factor G (EF-G) is responsible for translocating tRNA and mRNA down the ribosome during the translation process. The *Synechocystis* genome contains three genes (*slr1463/fusA*, *sll1098/fusB*, and *sll0830*) that encode different homologues of EF-G (*17*). EF-G has been identified as primary target within the translational system that becomes inactivated by reactive oxygen species (ROS) during photoinhibition (*17*). Oxidation of EF-G inhibits the *de novo* synthesis of the D1 protein, which in turn inhibits the repair of photodamaged photosystem II (PSII) (*17, 18*). The expression of redox-insensitive EF-G1 (encoded by *fusA*) by substituting a cysteine residue with serine has been shown to enhance the repair of PSII by accelerating the synthesis of the D1 protein under strong light (*19*). R461 of EF-G2 (encoded by *fusB*) is highly conserved across the green lineage. During ALE, it was found to be substituted to C, resulting in enhanced tolerance to HL stress (*9*). The mechanism by which a redox-sensitive cysteine in EF-G2_R461C_ increases its tolerance to oxidative damage is not yet clear. This study aimed to elucidate the underlying mechanisms of HL tolerance conferred by these mutations through transcriptomic and proteomic analysis of single and double mutant strains. The results provided insights into the functional components involved in HL tolerance and identified common and unique mechanisms across the mutated strains.

## Results

### HL tolerance of the evolved strain UMMM2 and two of its single mutations

The previous HL-ALE experiment (*9*) revealed that the EF-G2_R461C_ mutation emerged after the second round of mutagenesis, appearing with high frequency in several strains, including UMMM2 (100% frequency) (**Fig. 1A**). The NdhF1_F124L_ mutation, initially present at very low frequency (0.04%) in the original laboratory-type (LT) strain, appeared at higher frequencies after the final mutagenesis in UMMM2 (0.19%) and UMMM4 (100%) (**Fig. 1A**). UMMM2 carries both mutations (*9*) along with seven other high-frequency (>80%) mutations, including non-synonymous point mutations in four genes and frameshifts in three genes.

**Fig. 1.**
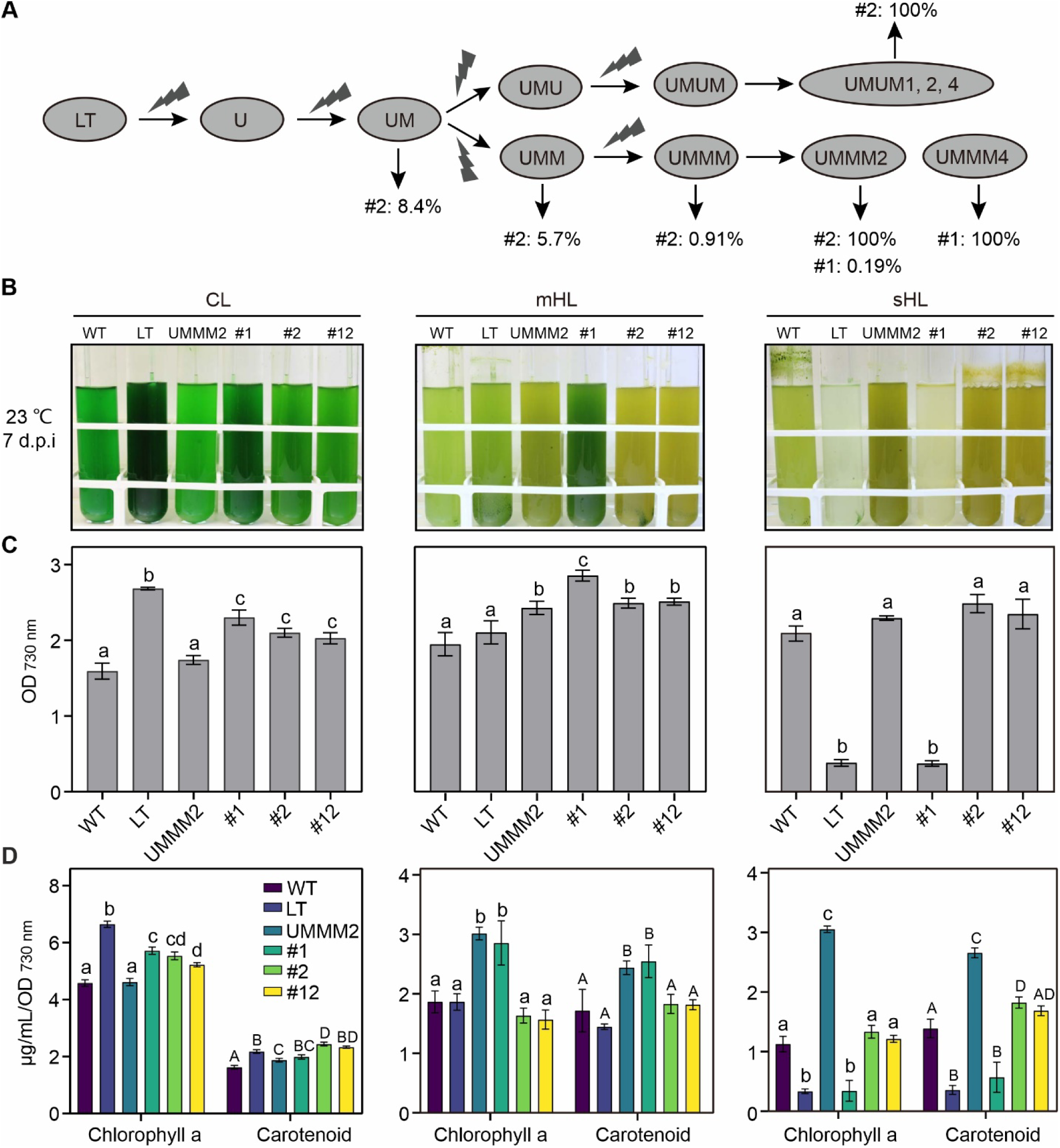
Origin and characteristics of the different strains under different light intensities. **(A)** The diagram depicts the mutagenesis process in the HL-ALE experiment, highlighting allele frequencies for two point mutations. U and M represent UV and methyl methanesulfonate (MMS) respectively, which were used as mutagens. Note that the UMMM2 strain harbored seven additional high-frequency (>80%) mutations, including non-synonymous point mutations in four genes (*rpoC1*, *gltA*, *slr0484*, and *ssr1480*) and frameshifts in three genes (*glcP*, *slr1546*, and *slr1975*). **(B)** Cultures of various strains (WT, LT, UMMM2, NdhF1_F124L_ (#1), EF-G2_R461C_ (#2), and NdhF1_F124L_+EF-G2_R461C_ (#12)) grown for a week under varying light intensities (CL, mHL, and sHL). Pre-cultures were grown for 7 days at CL at 23°C. The WT strain retains motility, while the LT strain (laboratory type) lacks it. All images were captured 7 days post-inoculation (d.p.i.). Notably, UMMM2 is the sole monoclonal strain from HL-ALE containing both NdhF1_F124L_ and EF-G2_R461C_ mutations. **(C)** Changes in final optical density (measured at 730 nm). Data points represent mean values with standard deviation, derived from three independent experiments. **(D)** Variations in chlorophyll and carotenoid content. Data points show mean values with standard deviation from three independent experiments. In **C** and **D**, statistical significance (p < 0.05) is indicated by different letters above error bars, as determined by one-way ANOVA with post-hoc Tukey HSD test.

Growth assays under various light conditions (CL, mHL, and sHL with 50, 700 and 1,200 μmol photons m^-2^ s^-1^, respectively) confirmed that NdhF1_F124L_ and EF-G2_R461C_ increase HL tolerance with a moderate CL trade-off (**Fig. 1B, C**). NdhF1_F124L_ cells showed superior growth at mHL but could not grow at sHL, while the EF-G2_R461C_ strain and the double mutant grew at sHL. The original wild-type motile strain from the Pasteur Collection (*20*) (WT) tolerated both HL conditions with a significant CL trade-off. The performance of UMMM2 was comparable to the point mutation strains under HL conditions but showed reduced growth and pigment content at CL **Fig. 1B-D**). Notably, UMMM2 survived at 2,000 μmol photons m^-2^ s^-1^, while EF-G2_R461C_ and the double mutant did not (*9*).

Taken together, NdhF1_F124L_ and EF-G2_R461C_ individually achieve growth rates under mHL and, in case of EF-G2_R461C_, sHL comparable to or higher than UMMM2. However, tolerance to even higher light intensities likely requires the additional mutations in UMMM2. The six strains (LT, WT, UMMM2, NdhF1_F124L_, EF-G2_R461C_, and NdhF1_F124L_+EF-G2_R461C_) serve as a model system for analyzing HL tolerance mechanisms evolved during ALE and their interplay in UMMM2.

### NdhF1_F124L_ and EF-G2_R461C_ are gain-of-function mutations

To explore the effects of the NdhF1 and EF-G2 mutations, insertional knockout (KO) and overexpression (OE) lines (using the strong *psbA2* promoter) were created (see Methods, **fig. S1A, B**). The NdhF1 KO strain (Δ*ndhf1*) showed slower growth than LT under CL and severe growth defects under mHL, indicating the importance of NdhF1 in HL acclimation (**Fig. 2A, B, fig. S1C**). Overexpression of NdhF1 (NdhF1-OE) or NdhF1_F124L_ (NdhF1*-OE) did not significantly affect growth compared to LT (**Fig. 2A, B; fig. S1C**). The Δ*ndhf1* strain exhibited increased Fv/Fm values relative to LT as reported (*21*), while the NdhF1_F124L_ strain showed intermediate values (**Fig. 2C**). CEF activity increased in NdhF1_F124L_ as described(*9*), but decreased in Δ*ndhf1* compared to LT (**Fig. 2D, E)**. Both OE strains rescued the Δ*ndhf1* growth defect and restored Fv/Fm and CEF activity to normal levels (**fig. S2)**. Similar results were obtained when NdhF1 and NdhF1_F124L_ were overexpressed in the Δ*ndhf1* background (**fig. S2**). These results, in particular the CEF activity data, suggest that NdhF1_F124L_ is a gain-of-function mutation, although overexpression alone was insufficient for HL tolerance.

**Fig. 2.**
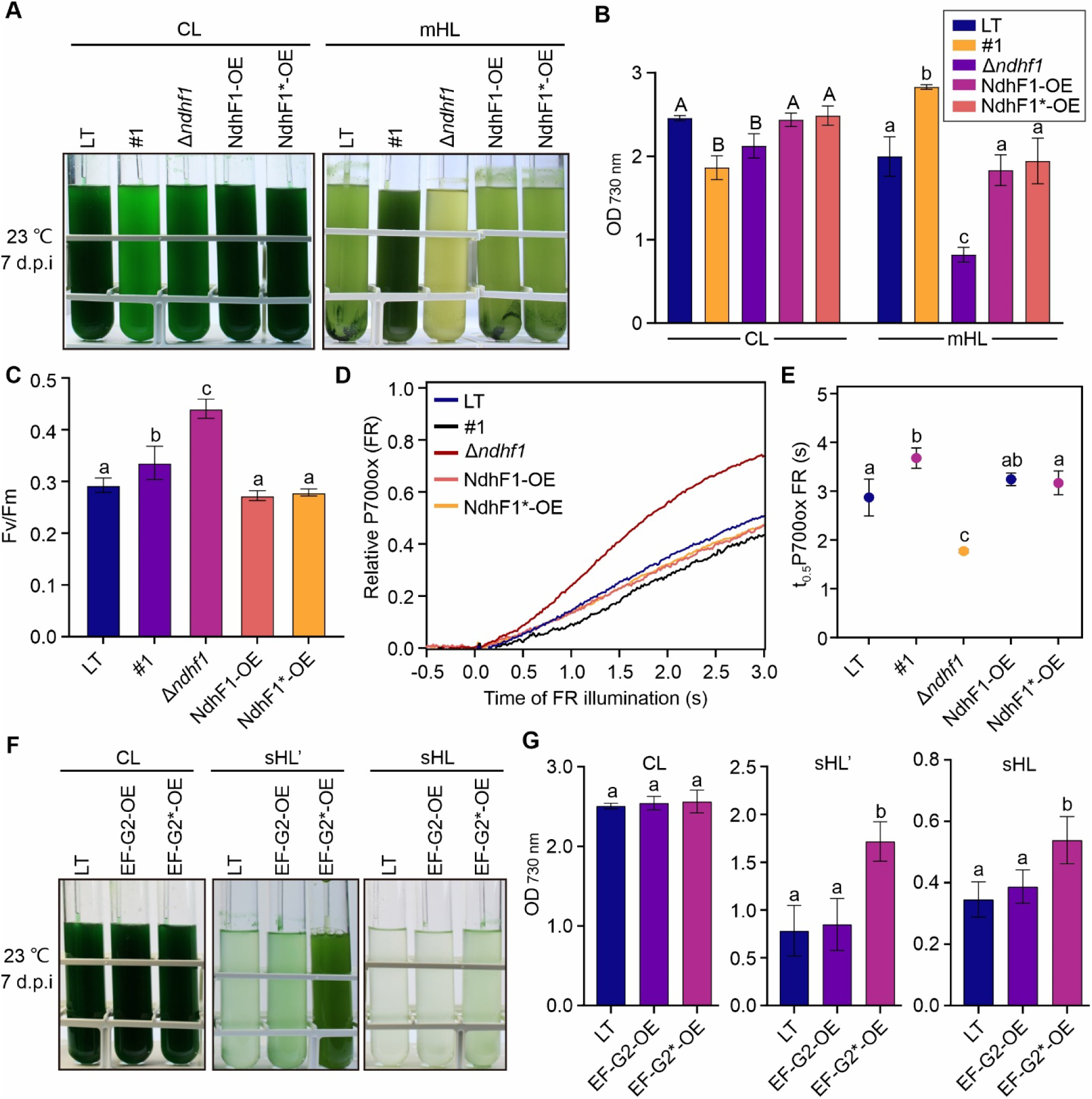
NdhF1_F124L_ and EF-G2_R461C_ are gain-of-function mutations. **(A)** Culture images of various strains (LT, NdhF1_F124L_ (#1), Δ*ndhf1*, NdhF1-OE, and NdhF1*-OE) grown for 7 days under CL and mHL conditions. Images, taken 7 d.p.i., are representative of three independent experiments. Strain descriptions: Δ*ndhf1* (*ndh*f1 knockout), NdhF1-OE (*ndhF1* overexpression), NdhF1*-OE (overexpression of *ndhFf1* with F124L mutation). **(B)** Final optical density changes (OD_730 nm_). Data shows mean ± SD from three independent experiments as in **A**. **(C)** Maximal PSII quantum yield (Fv/Fm) for LT, #1, Δ*ndhf1*, NdhF1-OE, and NdhF1*-OE cells. Mean ± SD from four independent experiments. **(D)** P700 oxidation kinetics for the same strains. Cells were exposed to far-red light (FR) to oxidize P700. Early P700 oxidation traces (ΔA820– 870) from four independent experiments are shown. Faster P700 oxidation (lower half-time values) indicates decreased CEF activity. **(E)** P700 oxidation half-times (t_0.5_ P700ox FR). Mean ± SD from the four experiments in **D**. **(F)** Images of LT, EF-G2-OE, and EF-G2*-OE cultures grown as in **A**. EF-G2-OE: *ef-g2/fusB* overexpression. EF-G2*-OE: overexpression of *ef-g2/fusB* with R461C mutation. sHL’: 1,000 μmol photons m^-2^ s^-1^. Images are representative of three independent experiments. **(G)** Final optical density changes (OD_730 nm_). Mean ± SD from the three experiments in **F**. For panels B, C, E, and G, different letters above error bars indicate statistically significant differences (p < 0.05) determined by one-way ANOVA with post-hoc Tukey HSD test.

For EF-G2, protein levels showed minor changes in EF-G2_R461C_ and NdhF1_F124L_+EF-G2_R461C_ strains under different light conditions, but a clear decrease in UMMM2 (**fig. S3A**), illustrating the (combined) effect of the seven additional high-frequency mutations (**Fig. 1A)**. We obtained a knockdown line (Δ*ef-g2/fusB*) but not a knockout line for EF-G2, which showed lower growth than LT under all conditions (**fig. S3B-E**). The OE strain of EF-G2_R461C_ (EF-G2*-OE) showed better growth than LT under sHL and sHL’ (1,000 μmol photons m^-2^ s^-1^), whereas the EF-G2-OE behaved similarly to LT (**Fig. 2F, G, fig. S3F**). Furthermore, when we transformed the KO construct into both OE strains, WT *ef-G2/fusB* copies could be completely deleted (**fig. S3B**), indicating that the EF-G2_R461C_ protein retains basic functions in protein translation and is a dominant gain-of-function mutation conferring HL tolerance.

### General trends in the transcriptomic HL response

RNA-seq analysis of the six strains under CL, mHL, and sHL conditions identified transcripts from 3,649 genes were identified across all 54 samples. Principal Component Analysis (PCA) revealed clear separation between the two HL conditions and CL, with most strains clustering closely under mHL and sHL (**Fig. 3A**), suggesting common cellular responses to HL. Differentially expressed genes (DEGs) were identified (see Materials and Methods), and strains with point mutations in LT showed more DEGs under HL conditions compared to LT, WT, and UMMM2 (**fig. S4A**). More DEGs were observed under CL and sHL conditions when comparing different strains to LT (**fig. S4B**), indicating substantial genotype-dependent differences in responses to these conditions.

**Fig. 3.**
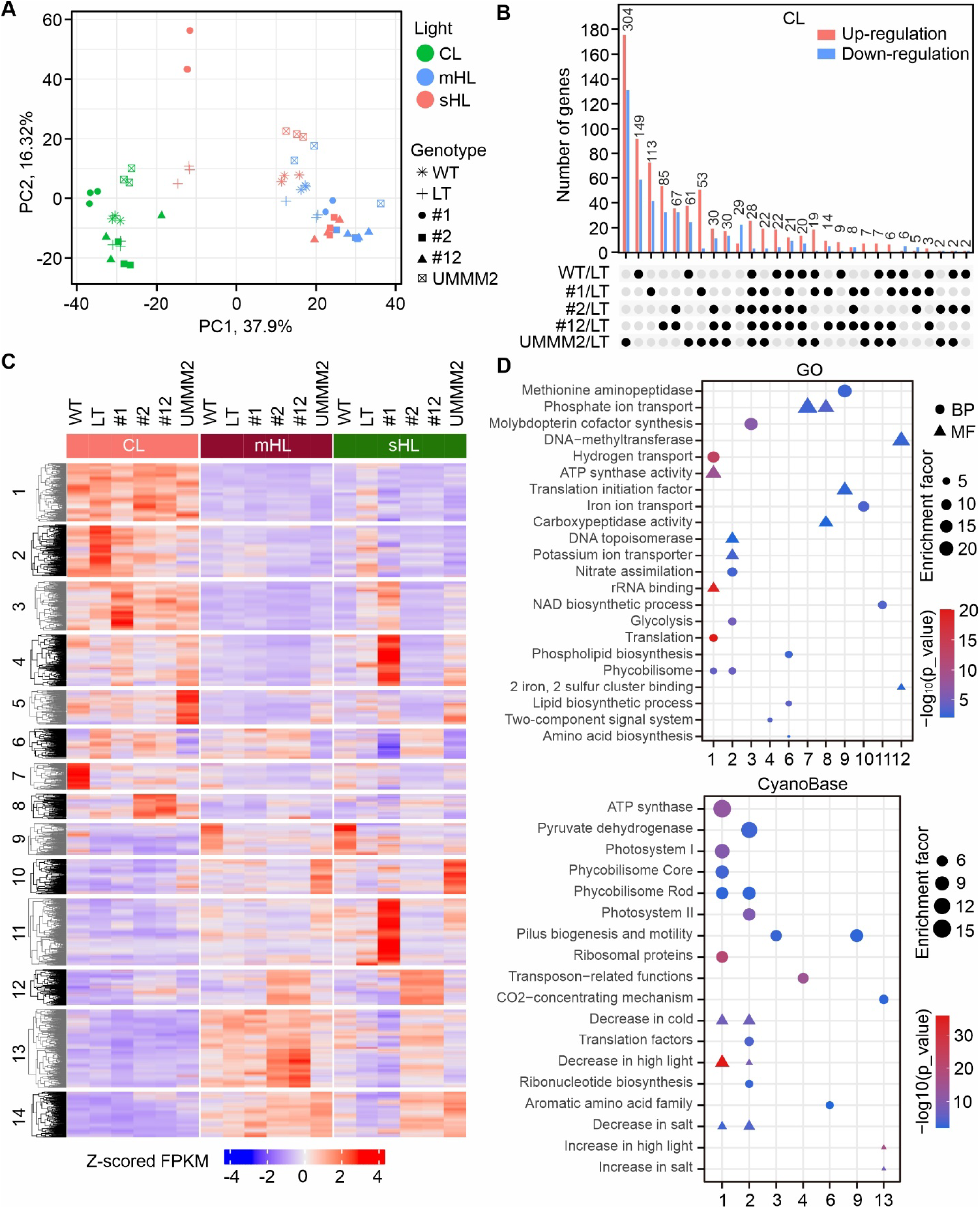
Overview and functional clustering analysis of transcriptome profiles. **(A)** Principal Component Analysis of CPM-normalized read counts for all identified transcripts across various genotypes and light conditions. The plot displays the two principal components with the highest variance percentages. One NdhF1_F124L_ (#1) sample at mHL (blue circle) was excluded due to its significant deviation from the other two replicates. Two NdhF1_F124L_ (#1) samples at sHL (red circle) were highly similar and appear as a single point on the plot. **(B)** UpSet plot illustrating the number of commonly up- or down-regulated Differentially Expressed Genes (DEGs) in various intersections of WT and four mutant strains (NdhF1_F124_ (#1), EF-G2_R461C_ (#2), NdhF1_F124_+EF-G2_R461C_ (#12), and UMMM2) compared to LT under CL conditions. The matrix below the x-axis shows intersections between the five genotypes relative to LT, with black dots indicating which genotypes are part of an intersection. The y-axis shows the number of up- or down-regulated transcripts in each intersection. Up- and down-regulated genes are analyzed separately but combined in the plot for easier visualization. Intersections are ordered by the total number of regulated genes (shown at the top of each set). **(C)** Heatmap of DEGs. Fourteen regulons were identified through hierarchical clustering of Z-score normalized FPKM values for 2,366 DEGs, using Euclidean distance and K-means algorithm. **(D)** Bubble plot showing non-redundantly enriched functions of different regulons. Functional terms were derived from the CyanoBase database and Gene Ontology (GO). Terms meeting the criteria of enrichment fold change > 2 and p-value < 0.01 (Fisher’s exact test) are displayed. Only regulons with found enrichment are shown. Symbol size indicates enrichment fold change. BP: biological process; MF: molecular function. Terms related to differential expression at HL, cold, and salt stresses (represented by triangles in the lower panel) were obtained from previous publications (*15, 29, 58–62*).

Pairwise overlap analysis revealed a large set of shared mHL/CL DEGs across all strains. WT, UMMM2, and NdhF1_F124L_ displayed the highest number of unique DEGs, while EF-G2_R461C_ shared most DEGs with NdhF1_F124L_+EF-G2_R461C_ (**Fig. 3b**; **fig. S4C-F**). UMMM2 showed more unique DEGs than shared ones with NdhF1_F124L_ or EF-G2_R461C_ (**Fig. 3b; fig. S4C-F**), suggesting its responses resulted from combined mutation effects rather than individual contributions.

Analysis of transcript accumulation for the nine mutated genes in UMMM2 revealed distinct patterns compared to other strains (**fig. S5**). In particular, reduced *fusB/ef-G2* mRNA goes along with the reduced levels of the EF-G2 protein in UMMM2 (**fig. S3A)**. This indicates that adaptive mutations may affect both protein activity and transcript accumulation of other mutated genes, adding complexity to the HL response in UMMM2.

In conclusion, the moderate HL response appears similar across genotypes at the transcriptome level. NdhF1_F124L_ and/or EF-G2_R461C_ effects represent only part of the response of UMMM2 response due to additional adaptive mutations. In UMMM2, adaptive mutations affect both protein sequences and mRNA expression of other mutated genes, indicating that HL adaptation alters gene expression patterns and regulatory loops.

### Common and genotype-specific pattern in the transcriptomic HL response

Analysis of 2,366 DEGs identified in genotype vs. LT or light conditions vs. CL comparisons revealed 14 regulons through hierarchical clustering (**Fig. 3C**). Functional enrichment analysis of these regulons provided insights into genotype-specific and light-dependent responses (**Fig. 3D**).

Under HL conditions, regulons 1-3 exhibited general down-regulation, with enrichment in photosynthetic functions (regulons 1 and 2) and translation-related processes (regulon 2) (**Fig. 3C, D; fig. S6**). This decrease in photosynthesis-related gene expression is characteristic of the HL stress response (*16*). However, the LT and NdhF1_F124L_ strains showed less severe decreases in PSII gene transcripts and notable up-regulation of the three *psbA* genes under sHL (**Fig. 3C, D; fig. S6**). Additionally, transcript accumulation for ribosomal subunits decreased under both HL conditions, except for LT and NdhF1_F124L_ at sHL (**fig. S6**), suggesting a slowdown in the translation process during HL.

Genotype-specific transcript patterns were observed in several regulons (**Fig. 3C, D**). NdhF1_F124L_ showed increased transcript accumulation in regulons 4 and 11 at sHL, enriched for “Transposon-related functions” and “NAD biosynthesis”. The WT strain displayed unique up-regulation in regulons 7 (enriched for ‘Phosphate ion transport’ and ‘Translation initiation factor’) and 9 (enriched for ‘Methionine aminopeptidase’ and ‘Pilus biogenesis and motility’) under different light conditions. Strains with EF-G2_R461C_ mutation showed distinct up-regulation in regulons 8 and 12, enriched for ‘Phosphate ion transport’ and ‘DNA methyltransferase’, respectively across all light conditions. In agreement with our other observations, the general transcript profile of the EF-G2_R461C_ and NdhF1_F124L_+EF-G2_R461C_ strains was highly similar. UMMM2 exhibited its own specific transcriptional responses in regulons 5, 6, and 10, consistent with its low overlap of DEGs with NdhF1_F124L_ or EF-G2_R461C_. Notably, regulon 10, enriched for ‘Iron ion transport’, showed significant up-regulation in UMMM2 under both HL conditions. Regulons 13 and 14, associated with ‘Increase in high light/salt’ responses, generally showed increased transcript abundance under HL conditions in all genotypes except LT and NdhF1_F124L_ at sHL.

These findings demonstrate genotype-specific and light-dependent control of regulon expression, suggesting distinct underlying responses contributing to HL tolerance.

### Gene sets specifically involved in the tolerance to strong HL

To identify DEGs directly involved in HL tolerance, we categorized strains into sHL-tolerant (WT, UMMM2, EF-G2_R461C_, NdhF1_F124L_+EF-G2_R461C_) and sHL-intolerant (LT, NdhF1_F124L_) groups. We then classified DEGs into clusters using fuzzy c-means algorithm (*22*) (**Fig. 4A-C; fig. S7**), focusing on clusters with expression patterns unique to sHL-tolerant strains. We analyzed WT and UMMM2 separately, comparing them to the two intolerant strains (LT and NdhF1_F124L_). Due to their similar responses, EF-G2_R461C_ and NdhF1_F124L_+EF-G2_R461C_ were analyzed together against the intolerant strains.

**Fig. 4.**
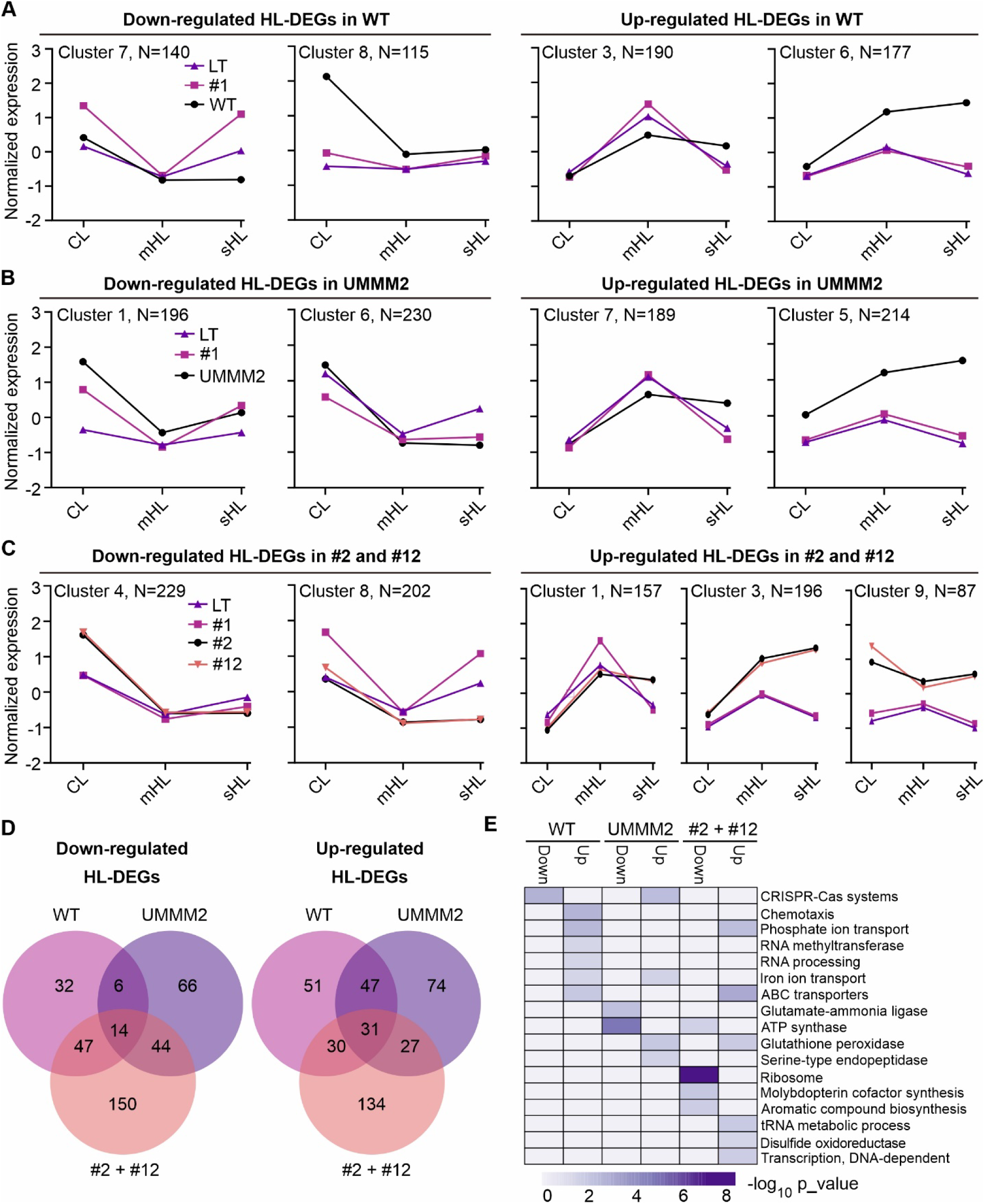
Identification of DEG sets associated with HL tolerance. (A-C) Strains are categorized into sHL-tolerant (WT, UMMM2, EF-G2_R461C_ (#2), and NdhF1_F124L_+EF-G2_R461C_ (#12)) and sHL-intolerant (LT and NdhF1_F124L_ (#1)) groups based on their ability to grow under sHL conditions. Line plots show averaged gene expression patterns identified by Fuzzy c-means soft clustering analysis for these groups (details in fig. S **7**). The plots highlight DEG groups with notably different regulation between sHL-tolerant WT (**A**), sHL-tolerant UMMM2 (**B**), and sHL-tolerant #2 and #12 (**C**), compared to the sHL-intolerant group (LT and #1). Gene numbers for each group are provided. The x-axis represents light intensities (CL, mHL, sHL), while the y-axis shows mean normalized expression. (**D)** Venn diagrams illustrating the number of shared and unique up- and down-regulated HL-tolerance related genes (HL-DEGs) in HL-tolerant strains: WT, UMMM2, and combined #2 + #12. (**E)** Heatmap displaying Gene Ontology (GO) terms enriched for up- and down-regulated HL-DEGs in WT, UMMM2, and combined #2 + #12 from **D**. The color scale represents -log_10_ transformed p-values from Fisher’s exact test.

For WT, we identified 8 distinct clusters (**fig. S7A**). Clusters 7 and 8 contained genes down-regulated in WT under both HL conditions, while intolerant strains showed up-regulation or similar expression (**Fig. 4A**, **fig. S7A**). These clusters may contain negative regulators of HL tolerance. Conversely, clusters 3 and 6 showed up-regulation in WT under HL conditions, suggesting positive regulators of HL tolerance (**Fig. 4A, fig. S7a**). After filtering with a membership value above 0.9, we identified 99 down-regulated and 159 up-regulated HL-tolerance-related DEGs (HL-DEGs) in WT.

Similar analyses for UMMM2 and EF-G2_R461C_/NdhF1_F124L_+EF-G2_R461C_ yielded 8 and 9 clusters, respectively (**Fig. 4B, C; fig. S7B, C**). We identified 130 down-regulated and 179 up-regulated HL-DEGs in UMMM2, and 255 down-regulated and 222 up-regulated HL-DEGs in EF-G2_R461C_/NdhF1_F124L_+EF-G2_R461C_.

Comparing HL-DEGs sets across sHL-tolerant strains revealed that 45% of down-regulated and 32% of up-regulated HL-DEGs from UMMM2 were shared with EF-G2_R461C_/NdhF1_F124L_+EF-G2_R461C_ (**Fig. 4D**), indicating the persistent impact of the EF-G2_R461C_ mutation in UMMM2. About 68% of HL-DEGs in WT were shared with UMMM2 or EF-G2_R461C_/NdhF1_F124L_+EF-G2_R461C_, but only a small subset was common to all three. Functional annotation based on GO term enrichment showed limited overlap of HL-DEGs among the three groups of strains, with no terms shared by all three (**table S1;** **Fig. 4E**). This suggests diverse regulatory mechanisms for HL acclimation, likely influenced by only a few point mutations.

### Common and genotype-specific control of proteomic responses to HL

To identify proteins associated with HL tolerance, we employed quantitative shotgun proteomics (*23*), using ^15^N-labeled whole-cell protein extracts of *Synechocystis* LT cells grown under CL and 200 μmol photons m^-2^ s^-1^ as an internal standard, which was spiked into each sample. Our analysis focused on 718 proteins detected across WT, LT, NdhF1_F124L_, EF-G2_R461C_, and NdhF1_F124L_+EF-G2_R461C_ (**Fig. 5A**), excluding the UMMM2 samples due to their high variation. PCA revealed a clear separation between HL and CL conditions, with WT, EF-G2_R461C_, and NdhF1_F124L_+EF-G2_R461C_ clustering closely under both mHL and sHL conditions (**Fig. 5B**).

**Fig. 5.**
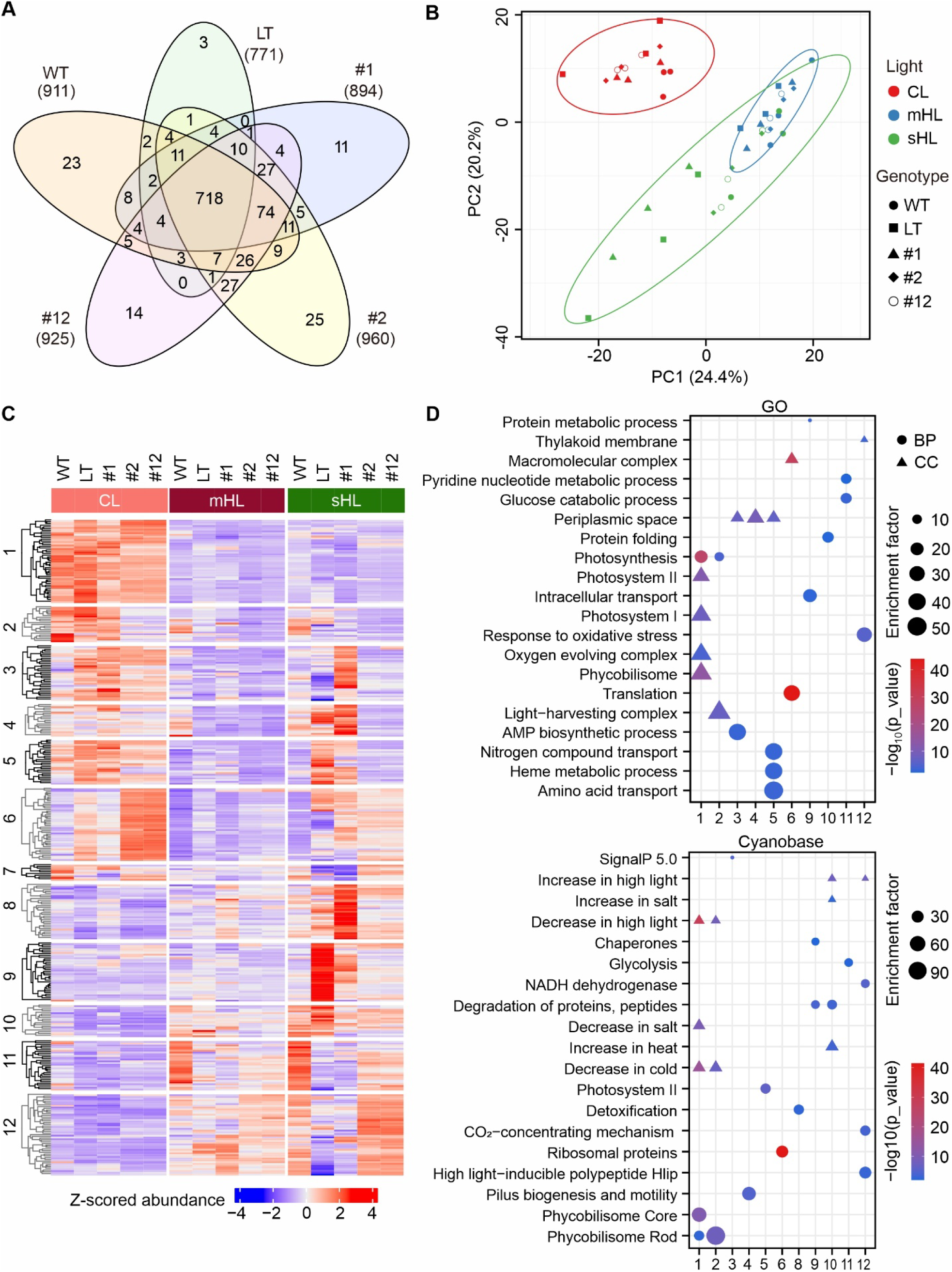
**Proteome changes in different strains under varying light intensities**. **(A)** Venn diagram showing the overlap and unique proteins accumulated in five strains: WT, LT, NdhF1_F124L_ (#1), EF-G2_R461C_ (#2), and NdhF1_F124L_+EF-G2_R461C_ (#12). Total proteins identified per strain are indicated in parentheses. **(B)** Principal Component Analysis of the relative abundance of 718 proteins common to all strains identified in **a**, across different genotypes and light conditions. The plot displays the two principal components with the highest variance percentages. **(C)** Heatmap of differentially expressed proteins (DEPs). Twelve distinct regulons were identified through hierarchical clustering of Z-scored relative abundance of 347 DEPs, using Euclidean distance and K-means algorithm. **(D)** Bubble plot showing enriched non-redundant functions of different regulons. The methodology used is identical to that described in Fig. 3D. BP: biological process; CC: cellular component.

We identified 347 differentially expressed proteins (DEPs) among the 718 common proteins (see **Methods**), grouped by hierarchical clustering into 12 distinct regulons. EF-G2_R461C_ and NdhF1_F124L_+EF-G2_R461C_ showed high similarity at the proteome level (**Fig. 5C**). Some regulons, particularly 5, 9, 11, and 12, displayed opposing expression patterns between sHL-tolerant and - intolerant strains at sHL (**Fig. 5C**), suggesting potential HL tolerance candidates.

Functional enrichment analysis revealed downregulation of photosynthesis-related proteins under HL conditions (regulons 1 and 2), and specific upregulation of translation and ribosomal proteins in EF-G2_R461C_-containing strains under CL (regulon 6) (**Fig. 5C, D; fig. S8**). Stress response proteins were strongly induced in LT and NdhF1_F124L_ at sHL (regulons 8 and 9). Metabolic processes and stress responses were upregulated in sHL-tolerant strains (regulons 11 and 12), including ’High light-inducible polypeptides (Hlips)’, ’CO_2_-concentrating mechanism (CCM)’, and ’NADH dehydrogenase’ (**Fig. 5D**). The gradual induction of HliC and HliD, components of the CO_2_-concentrating mechanism (SbtA, CmpA and CcmA), and ten NDH-1 complex subunits was observed in the three sHL-tolerant strains (**fig. S8**). Notably, the sHL-intolerant NdhF1_F124L_ strain showed the highest levels of NDH-1 subunits under mHL conditions (**fig. S8**), as previously observed by immunoblot analysis for NdhK (*9*).

In sum, our analysis suggests that the NdhF1_F124L_ mutation promotes increased accumulation of the NDH-1 complex at mHL, explaining its superior growth (see **Fig. 1**) and higher CEF activity (*9*) at this light intensity. The EF-G2_R461C_ mutation is associated with specific downregulation of phycobilisome rod proteins under all light conditions (regulon 2), a trait previously linked to increased HL tolerance (*24, 25*). Additionally, EF-G2_R461C_ increases ribosomal proteins under CL, and Hlip proteins, CO_2_ concentration proteins and NDH-1 subunits with increasing light intensity. In conclusion, the level of NDH-1, and consequently CEF, plays a crucial role in the HL tolerance acquired by adaptation, with increased NDH-1 levels correlating well with superior growth under HL conditions.

### Proteins specifically involved in the tolerance to strong HL

To identify key proteins involved in HL tolerance, termed HL-DEPs, we employed the fuzzy c-means algorithm to cluster all DEPs based on their expression profiles. We then identified proteins potentially suppressing or promoting HL tolerance by comparing their distinct expression profiles between sHL-tolerant and -intolerant strains.

Comparison of expression patterns between DEPs from WT and EF-G2_R461C_/NdhF1_F124L_+EF-G2_R461C_ with sHL-intolerant strains LT and NdhF1_F124L_ yielded five and seven distinct clusters, respectively (**fig. S9A, B**). Using the same approach as for HL-DEGs, we identified DEPs specifically down- and up-regulated in the WT in clusters 5 and 3 (**Fig. 6A**). For EF-G2_R461C_/NdhF1_F124L_+EF-G2_R461C_, we found two clusters (2 and 7) with specific down-regulation as light intensity increased, and cluster 3 featuring DEPs with specific up-regulation compared to the two mHL-intolerant strains (**Fig. 6B**).

**Fig 6.**
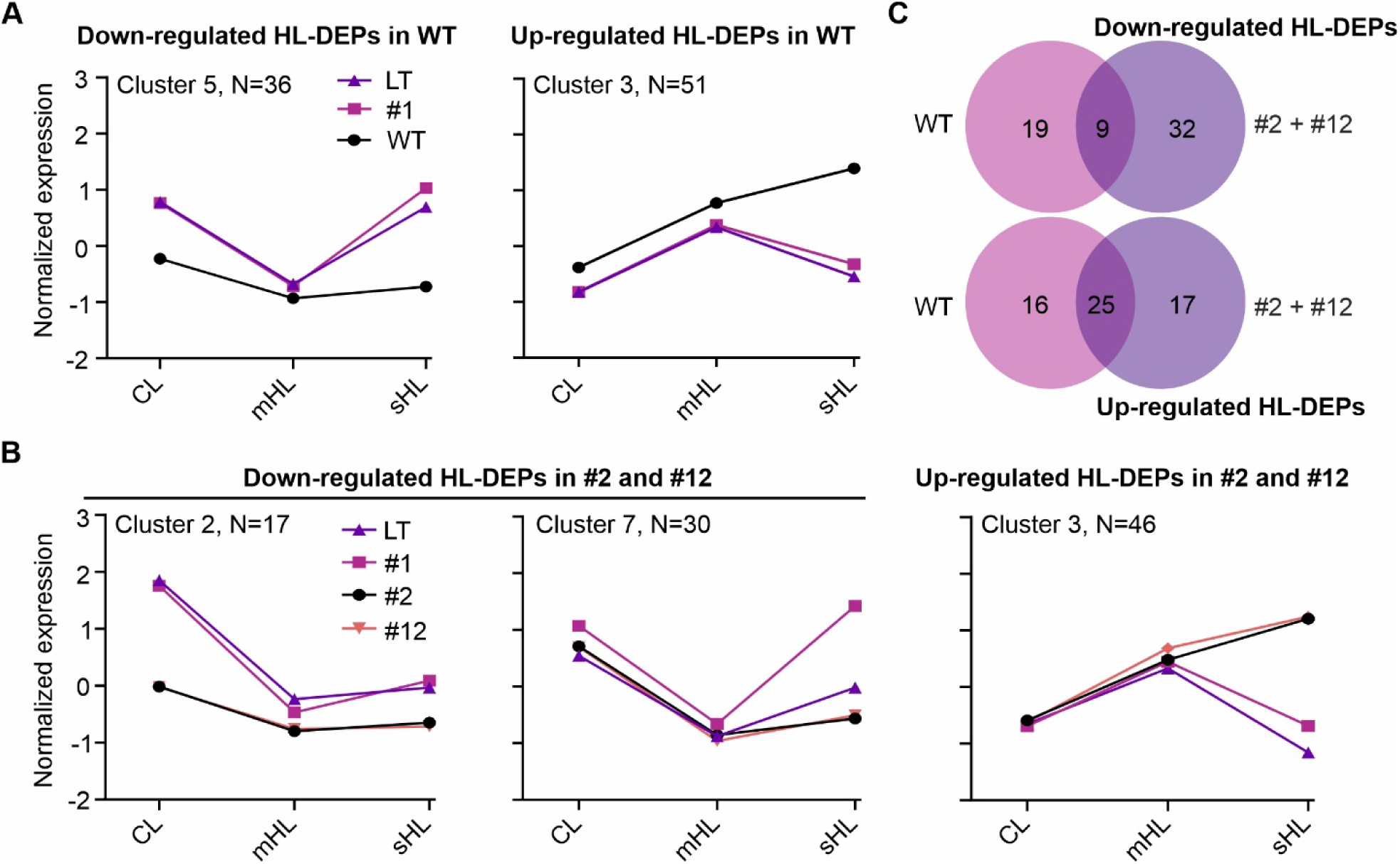
Identification of DEP sets associated with HL acclimation. (A, **B)** Line plots depicting averaged protein expression patterns among sHL-tolerant and intolerant strains. The plots highlight DEP groups showing notably different regulation between: sHL-tolerant WT (**B**), and sHL-tolerant EF-G2_R461C_ (#2) and NdhF1_F124L_+EF-G2_R461C_ (#12) (**B**) compared to the sHL-intolerant group (LT and NdhF1_F124L_ (#1)). See fig. S9 for details. The number of proteins in each group is provided. The x-axis represents light intensities (CL, mHL, sHL), while the y-axis shows mean normalized protein abundance. **(C)** Venn diagrams illustrating the number of shared and unique up- and down-regulated HL-tolerance related proteins (HL-DEPs) in HL-tolerant strains: WT and combined #2 + #12.

Applying a membership value threshold above 0.6 revealed 28 down-regulated and 41 up-regulated HL-DEPs in WT, while in EF-G2_R461C_ / NdhF1_F124L_+EF-G2_R461C_, we detected 41 down-regulated and 42 up-regulated HL-DEPs. PBS rod subunits (CpcA, B, C1, and C2) were identified as down-regulated HL-DEPs, while HliC, SbtA, CmpA, CcmA, and NDH-1 subunits were found as up-regulated HL-DEPs in EF-G2_R461C_/NdhF1_F124L_+EF-G2_R461C_.

Comparison of HL-DEPs between WT and EF-G2_R461_ /NdhF1_F124L_+EF-G2_R461C_ showed nine shared down-regulated and twenty-five shared up-regulated HL-DEPs (**Fig. 6C; table S2**). Notably, PBS rod proteins were among down-regulated HL-DEPs only in EF-G2_R461C_/NdhF1_F124L_+EF-G2_R461C_, while HliC, SbtA, and CcmA were common up-regulated HL-DEPs in both strains (**table S2**).

### Correlation of transcriptome and proteome changes

To investigate the correlation between transcript and protein changes, we conducted a global comparison of transcriptomic and proteomic datasets. After filtering with an ANOVA test (p<0.05), 660 high-confidence quantified proteins with corresponding transcripts were analyzed. Comparing fold changes of transcripts and proteins in mutants relative to LT at different light conditions revealed modest positive correlations, with Pearson correlation coefficients ranging from 0.25 (NdhF1_F124L_ vs. LT at sHL) to 0.56 (WT vs. LT at mHL). WT vs. LT consistently showed the highest coefficient at each light condition (**Fig. 7A; fig. S10A**), indicating that protein-level changes are only partially predictable from transcript-level changes when comparing genotypes.

**Fig. 7.**
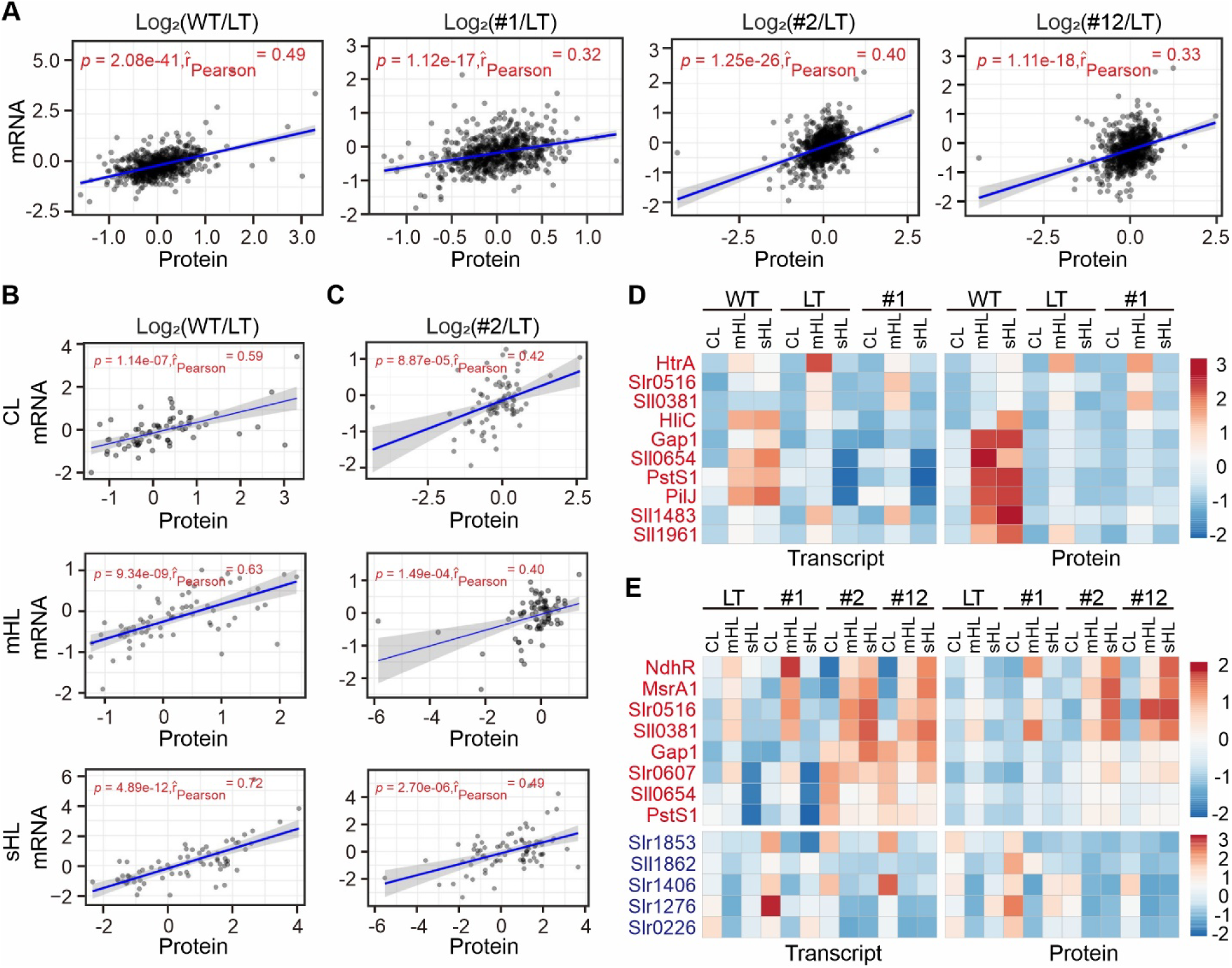
Comparison of transcriptome and proteome responses to different light intensities. **(A)** Scatter plots showing correlations between expression changes of all transcript-protein pairs at CL. The x- and y-axes represent log_2_-transformed protein and transcript ratios of WT, NdhF1_F124L_ (#1), EF-G2_R461C_ (#2), and NdhF1_F124L_+EF-G2_R461C_ (#12) relative to LT. Transcript-protein pairs are depicted as gray dots, with best-fit lines in blue. Pearson correlation coefficients (r) and student t-test p-values are provided. **(B, C)** Scatter plots illustrating correlations between expression changes of HL-DEPs and their corresponding transcripts in WT (**B**) and #2 (**C**). Log_2_-transformed protein and transcript ratios relative to LT under three light conditions (CL, mHL, and sHL) were used for correlation calculations. **(D, E)** Heatmaps showing abundance changes of transcript-protein pairs shared between HL-DEGs and HL-DEPs in sHL-tolerant WT (**D**) and sHL-tolerant #2 and #12 (**E**) compared to sHL-intolerant strains (LT and #1) under three light conditions (CL, mHL, and sHL). Only up-regulated shared HL-DEGs/DEPs were identified in WT. Genes/proteins from shared up-regulated and down-regulated HL-DEGs/DEPs groups are written in red and blue, respectively. The color bar indicates Z-score normalized FPKM for transcripts and relative abundance for proteins.

Higher correlations were observed when comparing transcript-protein pair fold changes from CL to mHL or sHL within the same genotype (**fig. S10B**), suggesting that proteome responses to HL are more predictable from transcriptome data when analyzing light condition effects in the same strain. An exception was the low correlation (0.20) for LT and NdhF1_F124L_ in the sHL vs. CL comparison, possibly due to cellular stress and growth arrest at sHL.

Analysis of WT HL-DEPs and EF-G2_R461C_ HL-DEPs showed higher correlation coefficients for WT vs. LT and EF-G2R461C vs. LT, respectively, across all light conditions compared to overall comparisons including all DEPs (**Fig. 7B, C**). Notably, only a small proportion of HL-DEPs had corresponding DEGs identified as HL-DEGs. For WT, none of the 28 down-regulated HL-DEPs had corresponding down-regulated HL-DEGs, while only 10 of 41 up-regulated HL-DEPs were identified as up-regulated HL-DEGs (**Fig. 7D**). Similarly, for EF-G2_R461C_, only 8 of 43 up-regulated HL-DEPs and 5 of 41 down-regulated HL-DEPs were identified as HL-DEGs (**Fig. 7E**). These findings suggest that only a fraction of the HL response involves regulation of protein accumulation through altered steady-state mRNA levels.

### Transcript-level independent regulation of CpcC2 accumulation may play a role in HL tolerance

The limited correlation between transcript and protein changes, along with the low overlap between HL-DEGs and HL-DEPs, suggests that factors beyond steady-state mRNA levels may influence HL-tolerance. Given that the EF-G2_R461C_ mutation affects an elongation factor, some protein accumulation changes in this strain might be directly caused by the mutation itself. To identify such cases, we searched for clear discrepancies between transcript and protein changes, focusing on anticorrelations. By plotting log_2_ (EF-G2_R461C_ vs. LT) values for transcript and protein changes across three light conditions, we identified 25 instances where proteins were at least 3-fold up- or down-regulated while corresponding transcripts were either oppositely or minimally regulated. Of these, 14 were HL-DEPs in EF-G2_R461C_ (**Fig. 8A**).

**Fig. 8.**
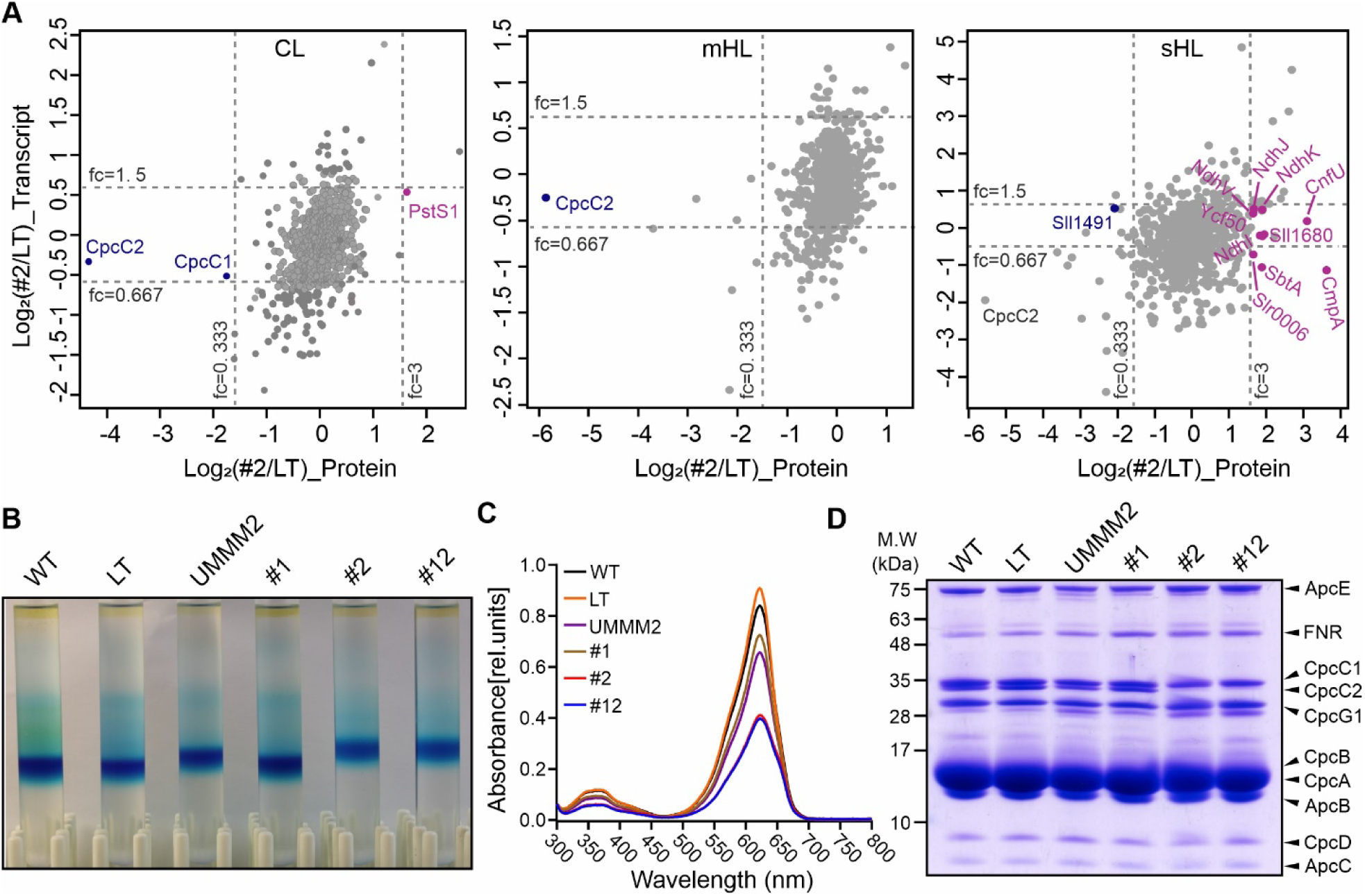
Transcript-independent regulation of CpcC2 contributes to HL tolerance in EF-G2_R461C_ strains. **(A)** Scatter plot comparing log_2_-transformed ratios of transcripts and proteins in EF-G2_R461C_ (#2) relative to LT (#2/LT) under three light conditions (CL, mHL, sHL). Highlighted are proteins with markedly different changes from their corresponding transcripts (at least 3-fold up- or down-regulated at protein level, while either oppositely or no more than 1.5-fold regulated at transcript level) across all light conditions. Among these, up-regulated and down-regulated HL-DEPs in #2 are labeled in pink and blue, respectively. **(B)** PBS size analysis using sucrose gradient. The blue band indicates purified PBS positions, with higher positions suggesting decreased PBS size. Strains: WT, LT, UMMM2, #1 (NdhF1_F124L_), #2 (EF-G2_R461C_), #12 (NdhF1_F124L_+EF-G2_R461C_). Image shown is representative of three independent experiments. **(C)** Absorption spectrum of purified PBSs at room temperature. Peak absorbance at 625 nm corresponds to phycocyanin (PC). Spectra normalized at 300 nm. Data represents means from three independent experiments. **(D)** Protein composition of purified PBSs. SDS-PAGE separation of proteins from purified PBSs, stained with Coomassie Brilliant Blue. Bands assigned to individual PBS proteins(*25, 51*). Gel shown is representative of three independent experiments as in **B**.

To validate these transcript-level-independent regulations in HL-tolerance, we investigated CpcC2, a down-regulated HL-DEP in EF-G2_R461C_ and NdhF1_F124L_+EF-G2_R461C_ strains. CpcC2 showed clear discrepancies between protein and transcript levels across all light conditions (**Fig. 8A**). In EF-G2_R461C_, CpcC2 protein levels were significantly reduced compared to LT, with only slight transcript decreases under various light conditions. CpcC2 is a phycobilisome (PBS) protein, part of the light-harvesting complex in cyanobacteria. PBS consists of an allophycocyanin core with six radiating rods, each composed of three phycocyanin hexamer discs connected by linker proteins, including CpcC2 for the distal disc (*26*). CpcC2 deletion results in smaller PBS size, which can lead to decreased photoinhibition and improved growth at higher light intensities (*25*). The drastic decrease of CpcC2 in EF-G2_R461C_ is expected to result in smaller PBS with less phycocyanin, consistent with previously observed reductions in 625 nm absorbance (*9*)(*9*)(*9*)(Dann *et al.*, 2021)(Dann *et al.*, 2021)(Dann *et al.*, 2021)(Dann *et al.*, 2021). To confirm this, PBSs were purified and analyzed by sucrose density gradient, showing that PBSs from EF-G2_R461C_ and NdhF1_F124L_+EF-G2_R461C_ shifted to a higher position, indicating smaller size (**Fig. 8B**). UMMM2 PBSs also showed a shift, albeit less pronounced. Absorption spectra of purified PBSs corroborated these findings (**Fig. 8C**). SDS-PAGE analysis of purified PBS subunits revealed that CpcC2 was almost undetectable in EF-G2_R461C_ and NdhF1_F124L_+EF-G2_R461C_, consistent with proteomics results (**Fig. 8D**). UMMM2 PBSs also showed a significant, though less drastic, decrease in CpcC2.

These results suggest that the EF-G2_R461C_ mutation leads to smaller antenna complexes due to CpcC2 linker protein down-regulation, occurring independently of steady-state transcript levels. This mechanism may contribute to the enhanced HL-tolerance observed in these strains.

### Overexpression of members of the phosphate regulon improves HL tolerance

From the small set of overlapping HL-DEGs and HL-DEPs showing correlation of transcript and protein levels (**Fig. 7D, E**), four up-regulated ones (Sll0381, Gap1, PstS1, and Sll0654) were shared between WT and EF-G2_R461C_. PstS1, a member of the ABC-type inorganic phosphate (Pi) transport system, significantly increases expression under Pi limitation stress (*27*). PstS1 and PstS2 bind Pi and are encoded by separate *pst* gene clusters (*27*). All *pst* genes showed increased transcript accumulation in both WT and EF-G2_R461C_ (**fig. S11A**), with ’Phosphate ion transport’ enriched among up-regulated HL-DEGs in both strains (**Fig. 4E**). Sll0654, a putative alkaline phosphatase, belongs to the phosphate (Pho) regulon, which is induced by Pi limitation and up-regulated under HL (*27, 28*). In the two sHL-intolerant LT and NdhF1_F124L_ strains, Pho regulon expression was lower and down-regulated under sHL (**fig. S11A**), suggesting a link between sHL tolerance and phosphate regulon expression.

PstS1 and Sll0654 showed increased or maintained expression at transcript and protein levels in WT and EF-G2_R461C_-carrying strains with increasing light intensities, contrasting with the sHL-intolerant strains. Since we have demonstrated transcriptional regulation of protein abundance for both PstS1 and Sll0654, a direct way to test their effect on HL tolerance is to overexpress them. Therefore, we generated overexpression (OE) lines PstS1-OE and Sll0654-OE (**fig. S11B**). Growth comparisons with LT under CL, mHL, and mHL’ (500 μmol photons m^-2^ s^-1^) showed that LT initially grew faster (**fig. S11C**). However, at mHL’ and mHL, LT growth retarded after 48h, while PstS1-OE and Sll0654-OE outperformed LT in final cell concentration and pigment content after 7 days (**Fig. 9A-C**, **fig. S11C**). Neither OE line grew at sHL, similar to LT (**Fig. 9**, **fig. S11C**).

**Fig. 9.**
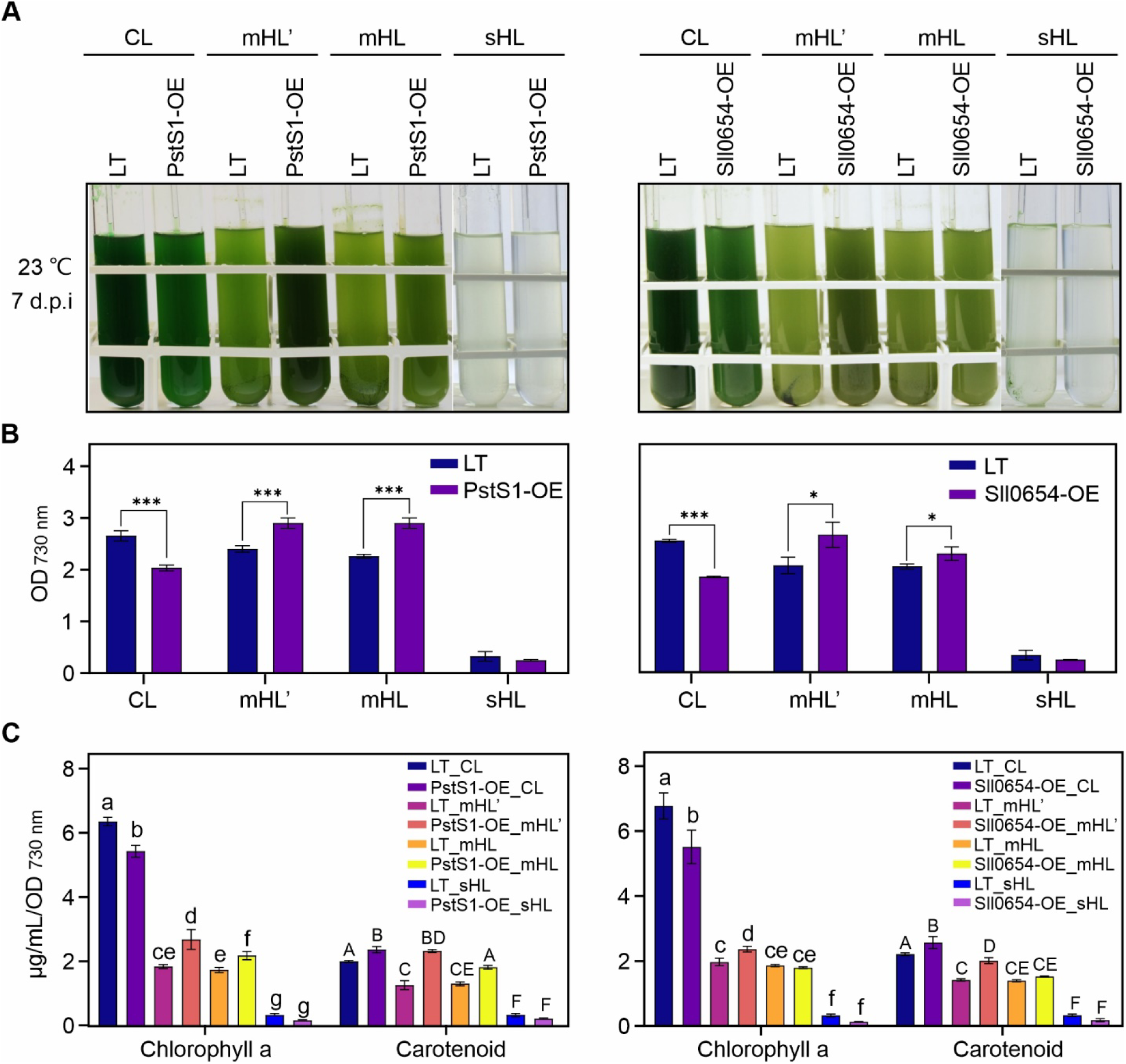
Overexpression of PstS1 and Sll0654 enhances high light tolerance. **(A)** Culture images of LT, PstS1-OE, and Sll0654-OE grown for 7 days under various light intensities (CL, mHL’, mHL, and sHL). mHL’ represents 500 μmol photons m^-2^ s^-1^. Pre-cultures were grown for 7 days at CL at 23°C. Images shown are representative of three independent experiments. (**B)** Optical density (OD_730 nm_) of LT, PstS1-OE, and Sll0654-OE cultures after 7 days of growth. Data shows mean ± SD from three independent experiments as in **a**. Statistical significance was determined using two-tailed Student’s t-test. *: p < 0.05, **: p < 0.01, ***: p < 0.001. (**C)** Changes in chlorophyll and carotenoid content. Data represents mean ± SD from three independent experiments as in **A**. Different letters above error bars indicate statistically significant differences (p < 0.05) as determined by one-way ANOVA with post-hoc Tukey HSD test.

These results suggest that increased Pi transport and metabolism is a conserved regulatory response for enhancing HL tolerance, albeit with a growth trade-off at low light conditions, as observed in WT, UMMM2, and EF-G2_R461C_ cells.

### Enhanced cyclic electron flow and decreased antenna size are common results of HL-ALE

Our findings indicated that NdhF1_F124L_ and EF-G2_R461C_ mutations enhance HL tolerance through increased CEF and reduced PBS size, respectively. To test if these regulations are common features of HL-ALE, we selected 9 representative lines from final HL-adapted monoclonal strains, along with UMMM2 from our previous ALE experiment (*9*). Only four of these 10 lines (UMUM1, UMUM4, UMMM2, and UMMM4) contained NdhF1_F124L_ and/or EF-G2_R461C_ (**Fig. 10A**).

**Fig. 10.**
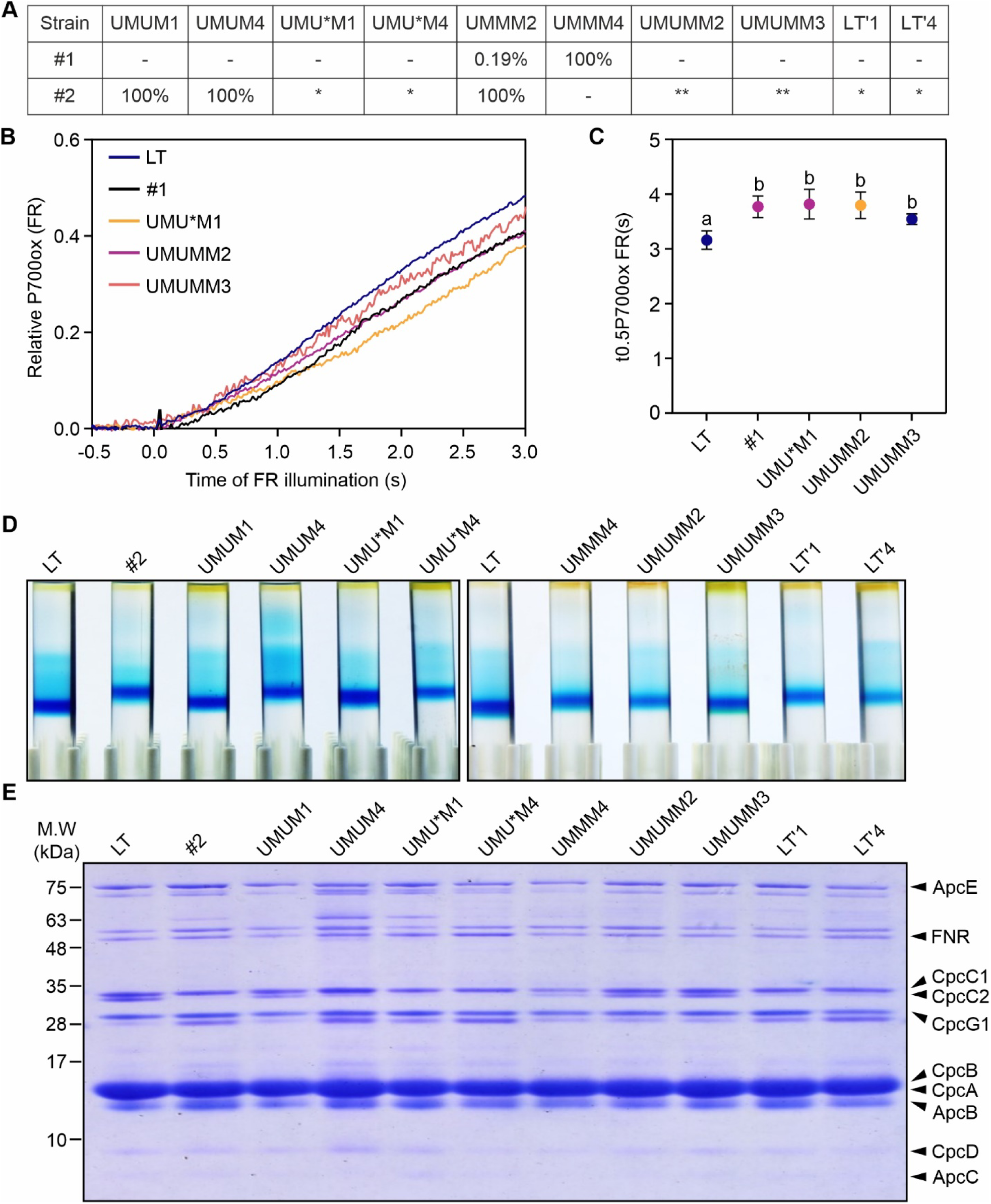
Enhanced cyclic electron flow and decreased phycobilisome size in HL-evolved strains. **(A)** Table showing representative HL-evolved strains selected from a previous HL-ALE experiment (*9*). Allele frequencies of both point mutations in these strains are also presented. Note that UMU*M and LT strains contain other *fusB* mutations at high frequency (*), whereas UMUMM strains contain such mutations at low/medium frequency (**). (**B)** P700 oxidation kinetics of LT, NdhF1_F124L_ (#1), UMU*M1, UMUMM2, and UMUMM3 cultures. Experimental settings are identical to those in Fig. 2E. Results for other strains can be found in **fig. S12**. (**C)** Half-times of P700 oxidation. Data shows mean ± SD from four independent experiments as in **B**. Different letters above error bars indicate statistically significant differences (p < 0.05) determined by one-way ANOVA with post-hoc Tukey HSD test. (**D)** PBS size analysis, following the same procedure described in Fig. 8B. (**E)** Protein composition of purified PBSs. Individual PBS proteins are assigned to bands as in Fig. 8D.

Analysis of P700 oxidation kinetics revealed that UMU*M1, UMUMM2, and UMUMM3 exhibited higher CEF activity than LT, comparable to NdhF1_F124L_ (**Fig. 10B, C**). However, UMUM4 and UMU*M4 showed significantly decreased CEF activity compared to LT (**fig. S12A, B**). The contrasting CEF activity levels between strains with the same evolution history, such as UMU*M1 and UMU*M4, suggest diverse HL-tolerance related mutations and regulations. Surprisingly, UMMM4 exhibited slightly lower CEF activity than LT (**fig. S12A, B**), despite possessing the F124L mutation in NdhF1 with 100% frequency, emphasizing that the effects of cooperating mutations cannot be reflected by a single mutation.

PBS purification and analysis revealed that most HL-adapted strains had reduced PBS size, with PBSs from UMUM4, UMU*M4, LT’1, and LT’4 shifting to a similar position as EF-G2_R461C_ (**Fig. 10D**). SDS-PAGE analysis of purified PBS subunit profiles showed that CpcC2 protein decreased to undetectable levels in these four strains with higher PBS shift, similar to EF-G2_R461C_ (**Fig. 10E**). This indicates that down-regulation of the CpcC2 linker protein is a common regulatory target in HL-ALE for achieving small PBS size, not solely through EF-G2_R461C_ but possibly by some other of the six mutant alleles of EF-G2 (**Fig. 10A**. However, UMUM1 exhibited similar trends in PBS shift and subunit patterns as LT (**Fig. 10D, E**), contrary to UMUM4, despite both possessing EF-G2_R461C_ with 100% frequency (**Fig. 1A, 10A**). This suggests that other mutations might mask the effects of EF-G2_R461C_ and confer HL-tolerance through different mechanisms.

In conclusion, a survey of representative HL-ALE strains demonstrated that increased CEF and decreased antenna size are recurring outcomes of HL-ALE, achievable through mutations in various gene sets beyond EF-G2_R461C_ and NdhF1_F124L_.

## Discussion

The development of HL tolerant strains through ALE has unveiled a complex genetic landscape with various combinations of point mutations (*9*). Achieving tolerance to HL conditions requires the synergistic effect of multiple genetic alterations, with mutations in different genes leading to similar epistatic outcomes. In strain UMMM2, mutations NdhF1_F124L_ and EF-G2_R461C_ co-occur, but NdhF1_F124L_ appears at a low frequency (see **Fig. 1**). When combined, these mutations show no additive effect on growth under various light conditions, with the negative effect of EF-G2_R461C_ on CEF prevailing (*9*). This lack of synergy suggests functional redundancy or slight antagonism in their effects on HL tolerance, explaining why they do not fully segregate together in ALE strains.

Proteomics analysis revealed a gradual increase in the expression of ten NDH-1 subunits from CL to HL conditions in sHL-tolerant strains, peaking at different light intensities for different strains (see **fig. S8**). The role of NdhF1 in HL adaptation is complex, with complete removal being detrimental under mHL conditions, while overexpression does not affect growth compared to LT at CL and HL (see **Fig. 2**). The NdhF1_F124L_ mutation showed downregulation of *ndhF1* transcripts under various light conditions (see **fig. S5**), suggesting a shift towards NDH-1MS complexes containing NdhF3, which may facilitate CEF during extended HL stress (*13, 14*). Clearly, NdhF1_F124L_ appears to be a gain-of-function mutation, conferring higher CEF than observed in either the *Δndhf1* KO or NdhF1 overexpressors (see **Fig. 2**). To fully understand the mechanisms at play, it remains to be determined whether the effects of mutating NdhF1 are specific or if similar results could be achieved by mutating NdhD1 proteins, which are present in NDH-1L but replaced by NdhD3 in NDH-1MS. Moreover, it is crucial to investigate whether the NdhF1 mutation affects the accumulation of NdhF1 itself, and how this mutation leads to increased overall accumulation of NDH proteins, potentially resulting in higher CEF.

Cyanobacteria typically downregulate photosynthetic complex subunits under HL to prevent excessive photodamage (*16, 29*), and a recent genome-wide CRISPR interference study revealed that PBS repression, while generally detrimental, is beneficial under HL (*30*). The most notable effect of the EF-G2_R461C_ mutation is the reduction in PBS size, primarily due to CpcC2 linker protein depletion, which mitigates excess light energy absorption (see **Fig. 8**). This adaptation aligns with findings in microalgae, where reducing the light-harvesting antenna cross-section improved growth under HL conditions (*24, 25, 31*). The EF-G2_R461C_ mutation induces an HL-like response even under CL conditions, resulting in reduced light energy input compared to other strains when exposed to strong HL. This adaptation, while protective under extreme HL, is not advantageous under lower light intensities due to limited light energy absorption, consistent with previous findings in a *Synechocystis* CpcC1C2-deficient strain (*25*).

Typically, intense illumination causes oxidative inactivation of elongation factors, suppressing protein synthesis and inhibiting photosystem II repair (*18, 19*). The suggestion that EF-G2_R461C_ mutation confers protection by adding a ROS-sensitive cysteine is controversial, given previous research showing increased photoprotection when replacing cysteine with serine in EF-G1 (*19*). Future studies should focus on examining de novo protein synthesis and PSII repair in EF-G2_R461C_ under sHL conditions.

HL conditions induce the expression of various proteins, including high light-inducible polypeptides (Hlips), CO_2_-concentrating mechanism (CCM) proteins, the Pho regulon, and NDH subunits (*12, 14–16, 27, 28, 32, 33*) primarily upregulates Hlips primarily upregulates and CCM proteins, with NdhF1 and CCM proteins, with NdhF1_F124L_ showing a lesser effect. The Pho regulon is induced specifically by EF-G2_R461C_. Maximum NDH-1 subunit induction is observed with NdhF1_F124L_ at mHL and EF-G2_R461C_ at sHL. Despite these varied responses, that individual adaptations are insufficient to confer sHL tolerance. This is evidenced by the sHL intolerance of NdhF1_F124L_ and our experiments overexpressing Pho regulon genes (see **Fig. 9**). The greater potency of EF-G2_R461C_ in conferring sHL tolerance suggests that a combination of multiple adaptive mechanisms is necessary for the observed HL tolerance in our evolved strains.

Examination of 10 ALE-derived HL-tolerant strains revealed that enhanced CEF and reduced antenna size are recurring characteristics (see **Fig. 10**). Three strains lacking the NdhF1_F124L_ mutation exhibited increased CEF, demonstrating that increased CEF can be achieved by mutations other than NdhF1_F124L_. Three strains without the EF-G2_R461C_ mutation showed reduced phycobilisome size linked to CpcC2 linker protein depletion. Therefore, downregulation of CpcC2 is a common regulatory target in HL-ALE, achieved through EF-G2_R461C_ and potentially other EF-G2 mutant alleles identified in the ALE strains (*9*). Since UMUM1 displayed a LT-like PBS size despite having EF-G2_R461C_ a 100% frequency (see **Fig. 10**), other mutations might mask EF-G2_R461C_ effects and confer tolerance through alternative mechanisms.

In sum, EF-G2 appears to be a potential hub for HL adaptation, being mutated in all HL-ALE strains (*9*). The mechanisms by which NdhF1 and EF-G2 mutations affect NDH complexes and CpcC2 accumulation remain unclear. HL-ALE produced strains with mutations in key proteins resulting in maximal tolerance. This adaptation strategy allows identification of unknown hubs for rewiring transcriptional and translational networks, facilitating the development of strategies to enhance photosynthetic efficiency.

## Materials and Methods

### *Synechocystis* strains and growth conditions

Various *Synechocystis* sp. PCC6803 strains were utilized, including the glucose-tolerant sHL-intolerant laboratory type (referred to LT), the original motile strain (referred to as WT), the HL-adapted monoclonal strain UMMM2 and corresponding strains with point mutations in the LT background as described previously (*9*). All cultures were grown at 23°C in Multi-Cultivator MC 1000-OD devices with an AC-700 cooling unit and warm white LED panel (Photon System Instruments, Brno, Czech Republic). Cells were pre-cultured in liquid BG11 cultures for about a week, then washed and inoculated at OD = 0.05 for photoautotrophic growth under different light conditions (CL, 50 μmol photons per m^-2^ s^-1^), moderate high light (mHL, CL, 700 μmol photons per m^-2^ s^-1^) or strong high light (sHL, CL, 1,200 μmol photons per m^-2^ s^-1^) for one week. Solid media growth used BG11 was supplemented with 0.75% (w/v) bacteriological agar (Carl Roth, Karlsruhe, Germany).

### Generation of *Synechocystis* insertional mutants and overexpressors

A non-replicative vector lacking the *sacB* selection cassette was employed to generate gene overexpression constructs, as previously described (*34*). Whole CDSs of candidate genes were amplified and inserted into the vector behind the *Synechocystis psbA2* promoter using Gibson Assembly® Cloning Kit (New England Biolabs, Ipswich, USA). The resulting plasmids, along with a chloramphenicol resistance (*CmR*) cassette, were inserted into the neutral site *slr0168* of the *Synechocystis* genome. Knockout constructs were made by flanking a spectinomycin resistance (*SpecR*) cassette with 500 bp upstream and downstream segments of target genes, then inserting this into a pUC57-derived vector.

Transformants were selected on BG11 plates with increasing concentrations of antibiotics (chloramphenicol from 5 to 20 μg/ml, spectinomycin from 20 to 50 μg/ml) and confirmed by genomic PCR.

### RNA isolation and transcriptome sequencing

For RNA extraction, 30 mL of cell cultures (OD_730_ ∼1.0) were harvested after one week of growth. Cells were processed with TRIzol (Invitrogen, Carlsbad, California, USA) at 65°C for 15 min, followed by RNA extraction using phenol-chloroform and isopropanol precipitation (overnight at -20°C). After washing with 75% ethanol, the precipitate was air dried and dissolved in RNase-free water. The RNA was treated with the TURBO DNA-*free*^TM^ kit (Invitrogen, MA, USA) to remove genomic DNA, and then purified using Direct-zol™ RNA MiniPrep Plus columns (Zymo Research, Irvine, California, USA). RNA integrity and quality were assessed using Nano Drop and the Agilent 2100 Bioanalyzer (Agilent, Santa Clara, Calif., USA). Only samples with an RNA Integrity Number (RIN) ≥ 6 were further processed. Ribosomal RNA depletion and RNA-Seq library generation were performed by Novogene Biotech (Beijing, China) using standard Illumina protocols. The RNA-Seq libraries were sequenced on an Illumina HiSeq 2500 system (Illumina, San Diego, Calif. USA) using a 150 bp paired-end sequencing strategy.

### Transcriptome data analysis

RNA-Seq reads were trimmed for adaptor sequences using *Trimmomatic* (*35*) and sequencing quality was assessed using *FastQC* (http://www.bioinformatics.babraham. ac.uk/projects/fastqc/). Paired-end reads were mapped to the *Synechocystis* sp. PCC6803 genome (GCF_000009725.1) using HISAT2 (*36*). Read counts were quantified using *featureCounts* (*37*). Principal component analysis (PCA) and UpSet plot were performed in the online web application iDEP (*38*) to determine the reproducibility between three biological replicates and to show group-wise comparisons of the overlapping genes, respectively. One sample (NdhF1_F124L_ at mHL) was widely separated from the other two replicates in the PCA and was omitted from the following analysis. Differentially expressed genes (DEGs) were obtained using *DESeq2* (*39*) by applying absolute log_2_ FC > 1 and an adjusted p-value < 0.05 in a contrasting group. FPKM (fragments per kilobase of transcript per million mapped reads) was calculated using the *fpkm* function in *DESeq2*. Genes with FPKM < 1 were determined to be low-expressed genes and removed from downstream analysis.

### Sample preparation for ^15^N-labelling based quantitative shotgun proteomics

The ^15^N-labelling labeled proteomics was performed as described before (*23*) with the following modifications. The *Synechocystis* LT strain was grown under constant illumination in BG11 medium containing 9 mM Na^15^NO_3_ as N source and subcultured continuously for 7 days to label the cells with ^15^N. The labeled cells were grown at CL and 200 μmol photons m^-2^ s^-1^ and then pooled together based on similar cell numbers to generate a reference standard. The ^15^N labelling efficacy of >98% was achieved.

The cell lysis and protein extractions were performed as reported previously (*40, 41*). The reference standard from ^15^N-labeled cells was spiked into the unlabeled samples from three biological replicates of each strain under different light conditions at a ^15^N/^14^N ratio of 0.8, based on protein content determined by the BCA protein assay kit (Thermo Fisher Scientific, MA, USA). Proteins were then precipitated in chloroform-methanol mixtures, as previously described (*42*). The pellets were dissolved in 6 M guanidine hydrochloride in 0.1 M HEPES (pH 8.5) containing protease inhibitors (Roche, Mannheim, Germany). The samples were incubated at 60°C for 10 min, sonicated three times for 20 sec, and clarified by centrifugation at 16,000 x g for 15 min. The clarified lysates were transferred to the filter units and digested according to the FASP method as described previously (*43*). The filters were centrifuged at 14,000 x g for 20 minutes and washed with 8 M urea in 0.1 M HEPES buffer (pH 8.5). Bound proteins were reduced by adding 10 mM DTT and incubated at 37°C for 10 min, followed by centrifugation at 14,000 x g for 20 min. Alkylation was performed by adding 0.05 M iodoacetamide for a 5-minute incubation. Samples were centrifuged as before, and filters were washed with 8 M urea in 0.1 M HEPES buffer (pH 8.5) and in HEPES buffer without urea. Proteins were digested with trypsin at 37°C for 16 hours. Digested peptides were purified with home-made C18 stage tips (*44*).

### Liquid chromatography and mass spectrometric analysis

Liquid chromatography-tandem mass spectrometry (LC-MS/MS) was conducted as previously described, with peptides separated over a 40-minute linear gradient of 5–80% (v/v) acetonitrile (ACN) (*42*).

### LC-MS/MS data analysis

Computational analysis of mass spectrometry (MS) measurements was performed using the public Galaxy platform (*45*). All used tools are available at https://usegalaxy.eu/ under the ‘Proteomics’ and ‘Convert Formats’ sections. The raw files in the proprietary Bruker format generated by the mass spectrometer were converted to the open mzML standard using the msconvert tool (*46*). Subsequent steps, including ion chromatogram extraction, identification and quantification of labeled (^15^N) and unlabeled (^14^N) peptides, as well as protein-level quantification, were conducted using the ProteomIQon tool chain (v0.0.7) (*47*).

Peptides were identified from MS/MS spectra by comparing them against the *Synechocystis* reference database employing a target-decoy strategy. Semi-supervised machine learning methods were utilized to integrate multiple search engine scores into a single consensus score (*48*). The results were statistically validated using q-value and PEP-value calculations, with thresholds set at 0.01 for q-values and 0.05 for PEP-values. Peptide abundance was determined by calculating the area under the curve of extracted ion chromatograms. Subsequently, quantified peptide ions were aggregated at the protein level.

Missing values were imputed using the K-Nearest Neighbors (KNN) method within the *imputeLCMD* package (*49*) at the protein level prior to statistical analysis among replicates. Proteins with high confidence quantitation were filtered by one-way ANOVA with p value <0.05. Then, differentially expressed proteins (DEPs) were determined by applying the fold change >2 in at least in one sample per genotype relative to LT in the three light conditions or in at least one sample in HL relative to CL in the different genotypes.

### Protein extraction and immunoblot analysis

*Synechocystis* cells were lysed in a buffer containing 0.4 M sucrose, 10 mM NaCl, 5 mM MgCl_2_ 20 mM Tricine (pH 7.9) and 0.5 mM PMSF with a pre-cooled TissueLyser II homogenizer (Qiagen, Venlo, Netherlands), and the insoluble debris was removed by centrifugation for 10 min at 10,000 rpm at 4 . Protein concentration was measured by BCA assay (Thermo Fisher Scientific, MA, USA). The whole cell lysate approximated equivalent to 20 μg protein content was loaded onto 10% (w/v) Tris-glycine SDS-PAGE. After gel running, separated proteins were transferred to PVDF membranes (Immobilon-P; Millipore, Burlington, MA, USA) with the Trans-Blot Turbo system (Bio-Rad, Hercules, CA, USA), and visualized by staining with Coomassie Brilliant Blue R-250 to keep equal loading. The membranes were blocked with 5% (w/v) milk in TBS-T buffer for 2 h at room temperature. After blocking, the membranes were incubated with the primary antibody (anti-EF-G2, 1/10,000; provided by Prof. Nishiyama) diluted in 3% (w/v) milk with TBS-T overnight at 4 .

Protein signals were detected by enhanced chemiluminescence using the SuperSignal™ West Pico PLUS Chemiluminescent Substrate (ThermoFisher Scientific, Waltham, MA, USA) and a Fusion FX ECL reader system (Vilber Lourmat, Collégien, France). Signal intensities were quantified using ImageJ software (National Institutes of Health).

### Determination of pigment content

After growth, 1 mL *Synechocystis* cells at OD_730 nm_ = 1.0 were centrifuged at 12,000 rpm for 1 min. The pellet was then resuspended in 1 mL 100% methanol and incubated overnight at 4 in the dark. The supernatant containing pigments was centrifuged at 4 for 5 min at 12,000 rpm and analyzed at 470, 665, 720 nm with a spectrophotometer (Ultrospec 2100 pro, Biochrom Ltd., Cambridge, UK). Quantification of chlorophyll and carotenoid content was performed as described before (*50*).

### Room temperature fluorescence measurement

For measurement of PSII fluorescence, *Synechocystis* cells were dropped onto BG11 agar plates at OD_730 nm_ of 10 and grow overnight. The PSII quantum yield parameters were determined with a FluorCam 800MF (Photon Systems Instruments, Brno, Czech Republic) as described previously (*9*) .

### P700 analysis

*Synechocystis* cells were cultured photoautotrophically in BG11 medium with an initial OD_730 nm_ of 0.05 in shaken flasks (140 rpm) at 23 °C under CL to exponential phase (OD_730 nm_ ∼0.8-1.0). Then the cells were harvested and treated for P700 oxidation analysis with DUAL-PAM-100 instrument (Walz GmbH, Germany), as described previously (*34*).

### Purification and analysis of phycobilisomes (PBSs)

PBSs were purified as described previously (*51*). Briefly, 100 mL cells in exponential phase (OD_730 nm_ ∼1.0) cultured at CL were harvested and washed twice with 0.75 M potassium phosphate (KP) buffer, pH 7.0. Cell pellets were resuspended in 1 ml KP buffer containing protease inhibitor cocktail (Roche, Mannheim, Germany). Cells were disrupted by vortexing with glass beads at 30 Hz for 5 min twice. The broken cell extracts were centrifuged at 4,000 rpm for 20 min, and the supernatants were collected and incubated with Triton X-100 (Carl Roth, Karlsruhe, Germany) at a final concentration of 2% (v/v) for 20 min at room temperature in the darks with gentle shaking. The unbroken cells and debris were removed by centrifugation at 20,000 rpm for 20 min at 15 °C. The dark blue supernatant was loaded onto a 10-40% (w/v) linear sucrose gradient in 0.75 M KP and centrifuged at 30,000 rpm for 16 h at 15 °C. After centrifugation, the blue bands were collected directly with a syringe, and then buffer-exchanged into 200 µL phosphate-buffered saline buffer, pH 7.0, using a 10 kDa Amicon filter. To measure the absorption spectra of purified PBSs, 10 µL PBS solution was mixed with 90 µL phosphate-buffered saline buffer in a 96-well microplate and recorded from 300 nm to 800 nm using a plate reader (Tecan, Maennedorf, Switzerland) at room temperature.

For subunit analysis, 20 µL of PBS solution was mixed with SDS loading buffer and incubated at 42 °C for 2 h. After incubation, the samples were run on an SDS-PAGE gel (12%) with 6M urea and visualized by Coomassie Brilliant Blue staining.

## Data availability

RNA-Seq data were deposited in the Gene Expression Omnibus repository with the accession number GSE268898. Proteomics data have been deposited with the ProteomeXchange consortium via the PRIDE (*52*) partner repository under accession number PXD052516. All additional data are available upon request from the corresponding author.

## Bioinformatic analysis and statistics

Heatmaps with hierarchical clustering were generated using *ComplexHeatmap* v2.10.0 packages (*53*) in R v4.1.0. Functional enrichment analysis was performed in Perseus v2.0.5.0 (*54*) with CyanoBase (http://genome.microbedb.jp/cyanobase/) and Gene Ontology (GO) terms (*55*). The significance of the enrichment was calculated with Fisher’s exact test. These terms with p value < 0.01 and enrichment fold change >2.0 are considered to be significantly enriched. The enrichment bubble plot was generated using *ggplot2* package (*56*) in R v4.1.0. The pairwise Pearson’s correlation coefficients between changes of transcript and protein were calculated by running the *ggscatterstats* function in R package *ggstatsplot* (https://github.com/IndrajeetPatil/ggstatsplot).

For fuzzy C-means clustering, the expression profiles of transcripts and proteins were grouped into different clusters using the fuzzy C-means algorithm with the *Mfuzz* package in R v4.1.0 (*57*). Before clustering, the FPKM of the transcripts and the relative abundance of the proteins were transformed using the *standardize* function in the *Mfuzz* package. The center of the expression change profile in each cluster was visualized by a line plot. A cluster membership value >0.9 or 0.6 was used as a threshold for filtering genes or proteins respectively with the best fit to the centralized expression pattern in each cluster.

GraphPad Prism version 10.1.0 software was used for graphs and statistical analyses using one-way ANOVA with post-hoc Tukey HSD test and two-tailed Student’s t-tests.

## Supplementary Materials This PDF file includes

Figs. S1 to S17 Tables S1 and S2

## Acknowledgements

We thank Prof. Nishiyama from Saitama University for providing the EF-G2 antibody.

## Author contributions

D.L. and W.C. conceived the project. D.L. provided funding. D.L. and W.C. designed all experiments. W.C. performed the experiments with support of M.D., C.O. and S.S. in the proteomics experiment. E.A.S. helped with the transcriptomics analysis and interpretation of the results. D.L. and W.C. wrote the manuscript. All authors read and approved the final manuscript.

## Data and materials availability

RNA-Seq data were deposited in the Gene Expression Omnibus repository with the accession number GSE268898. Proteomics data have been deposited with the ProteomeXchange consortium via the PRIDE (*52*) partner repository under accession number PXD052516. All additional data and materials are available upon request from the corresponding author.

