## Supplementary for "High-light adaptation in *Synechocystis* by accumulating NDH proteins and depleting specific phycobilisome linker proteins"

Weiyang Chen *et al.*

**This PDF file includes:**

Supplementary Text
Figs. S1 to S12
Tables S1 to S2

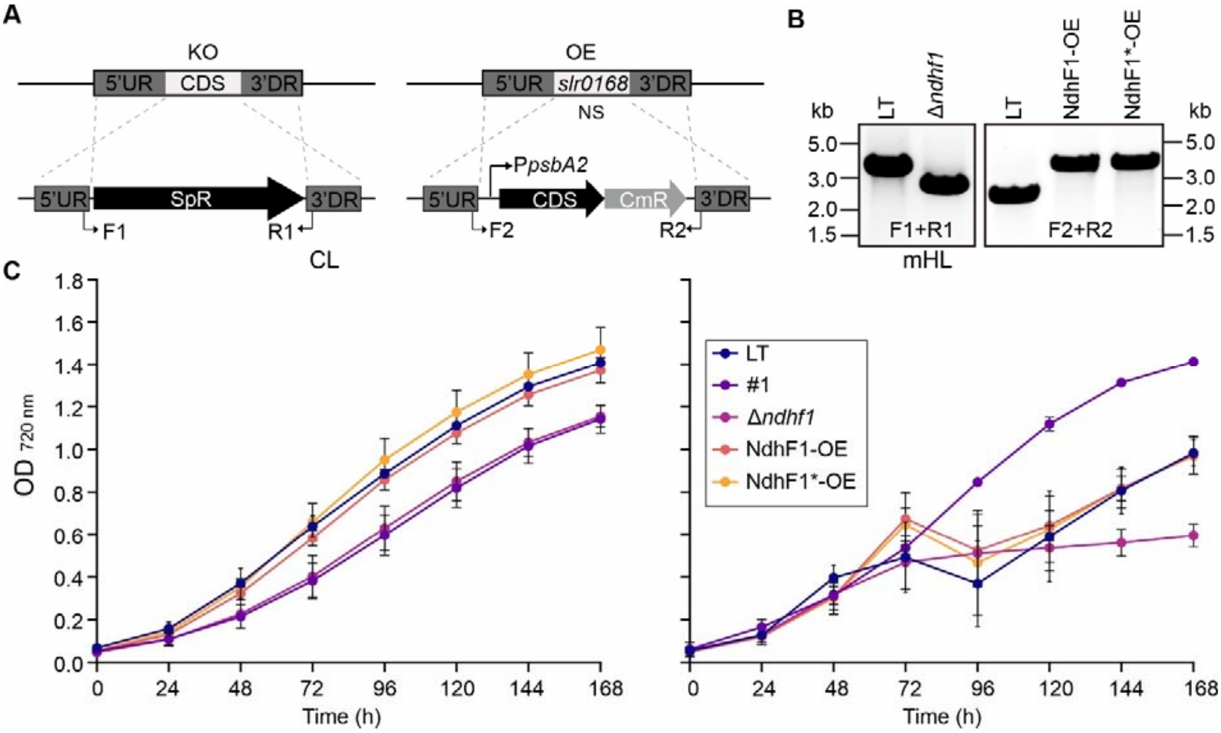


**Fig. S1.**

**Generation of *ndhF1* knockout and overexpression strains and growth analysis. (A)** Outline of the strategy for gene knockout (KO) and overexpression (OE) in *Synechocystis*. For KO, the target gene was substituted with a spectinomycin resistance gene cassette (*SpR*). For OE, the target gene, controlled by the *psbA2* promoter ($P_{psbA2}$) and linked to a chloramphenicol resistance gene cassette (*CmR*), was inserted into a neutral site (*slr0168*) of the *Synechocystis* LT genome. The diagram indicates the positions of the forward (F1, F2) and reverse (R1, R2) primers used for segregation control. **(B)** PCR analysis confirmed complete segregation of the KO and OE strains. *Δndhf1* represents the KO of *ndhf1*, NdhF1-OE denotes the OE of *ndhF1*, and NdhF1*-OE indicates the OE of *ndhF1* with F124L mutation. **(C)** Growth curves of LT, #1, *Δndhf1*, NdhF1-OE, and NdhF1*-OE under CL and mHL conditions. Optical density ($OD_{720\ nm}$) was measured hourly using the built-in real-time monitor system in Multi-Cultivator OD-1000 devices. For clarity, values at 24-hour intervals were plotted. The data shown represents the mean ± SD from three independent experiments, as in **Fig. 2A**.

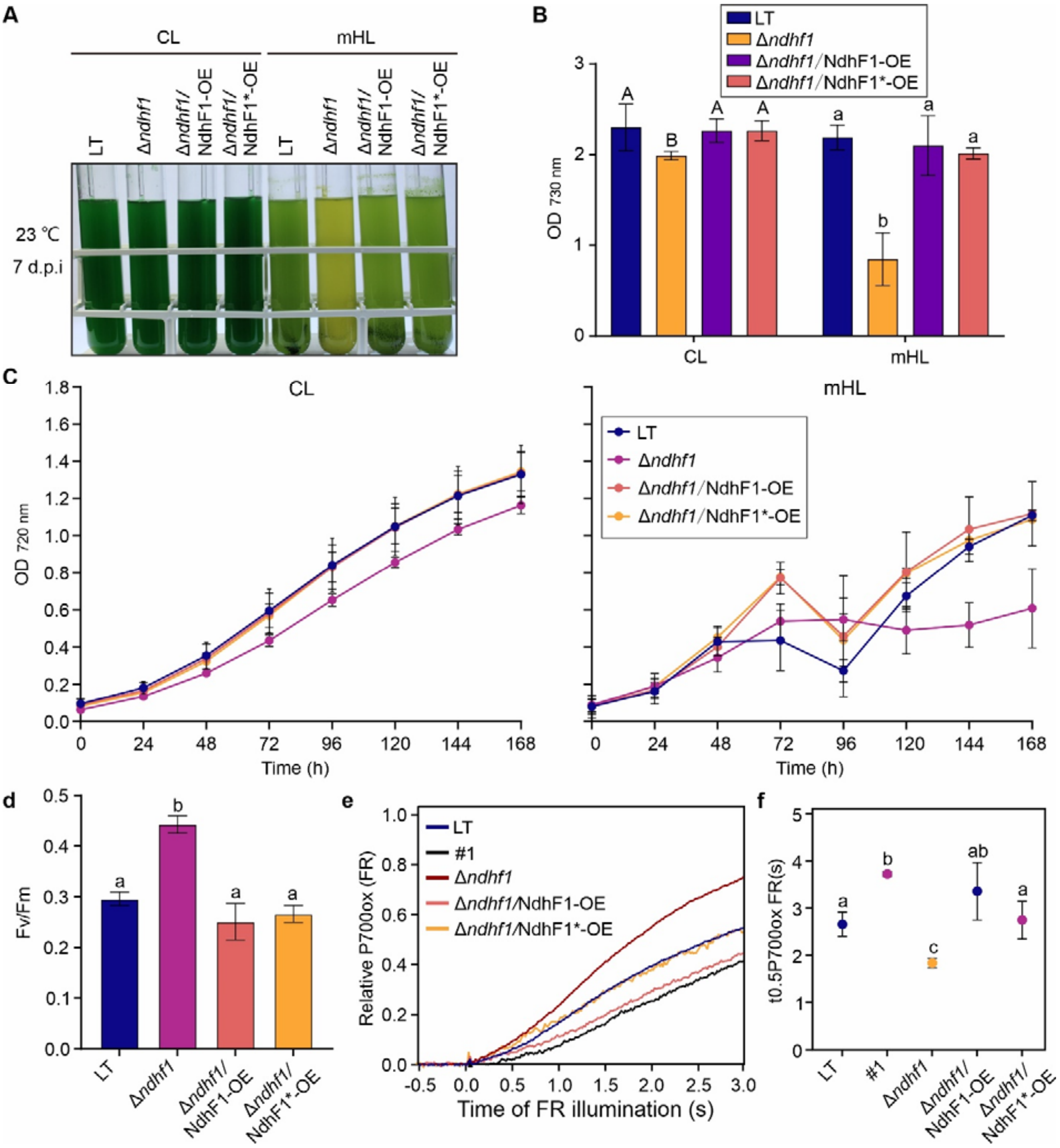


**Fig. S2.**

**Characterisation of *ndhF1* overexpressors in the *Δndhf1* background. (A)** LT, *Δndhf1*, *Δndhf1*/NdhF1-OE, and *Δndhf1*/NdhF1*-OE cultures after 7 days of growth under CL and mHL conditions. Pre-cultures were grown for 7 days at CL at 23°C. The image shown is representative of three independent experiments. *Δndhf1*/NdhF1-OE and *Δndhf1*/NdhF1*-OE represent KO of *ndhF1* in NdhF1-OE or NdhF1*-OE backgrounds, respectively. **(B)** Changes in final optical density ($OD_{730\ nm}$). The data shows the mean ± SD from three independent experiments, as in **a**. Different letters above error bars indicate statistically significant differences ($p < 0.05$), determined by one-way ANOVA with post-hoc Tukey HSD test. **(C)** Growth curves of LT, *Δndhf1*, *Δndhf1*/NdhF1-OE, and *Δndhf1*/NdhF1*-OE under CL and mHL conditions. The data represents the mean ± SD from three independent experiments, as in **A**. **(D)** Maximal PSII quantum yield (Fv/Fm) for LT, #1, *Δndhf1*, *Δndhf1*/NdhF1-OE, and *Δndhf1*/NdhF1*-OE cells. Mean ± SD from four independent experiments. **(E)** P700 oxidation kinetics for the strains in panel **A**. Four independent experiments were conducted. **(F)** P700 oxidation half-times. Mean ± SD from the four experiments in **E**.

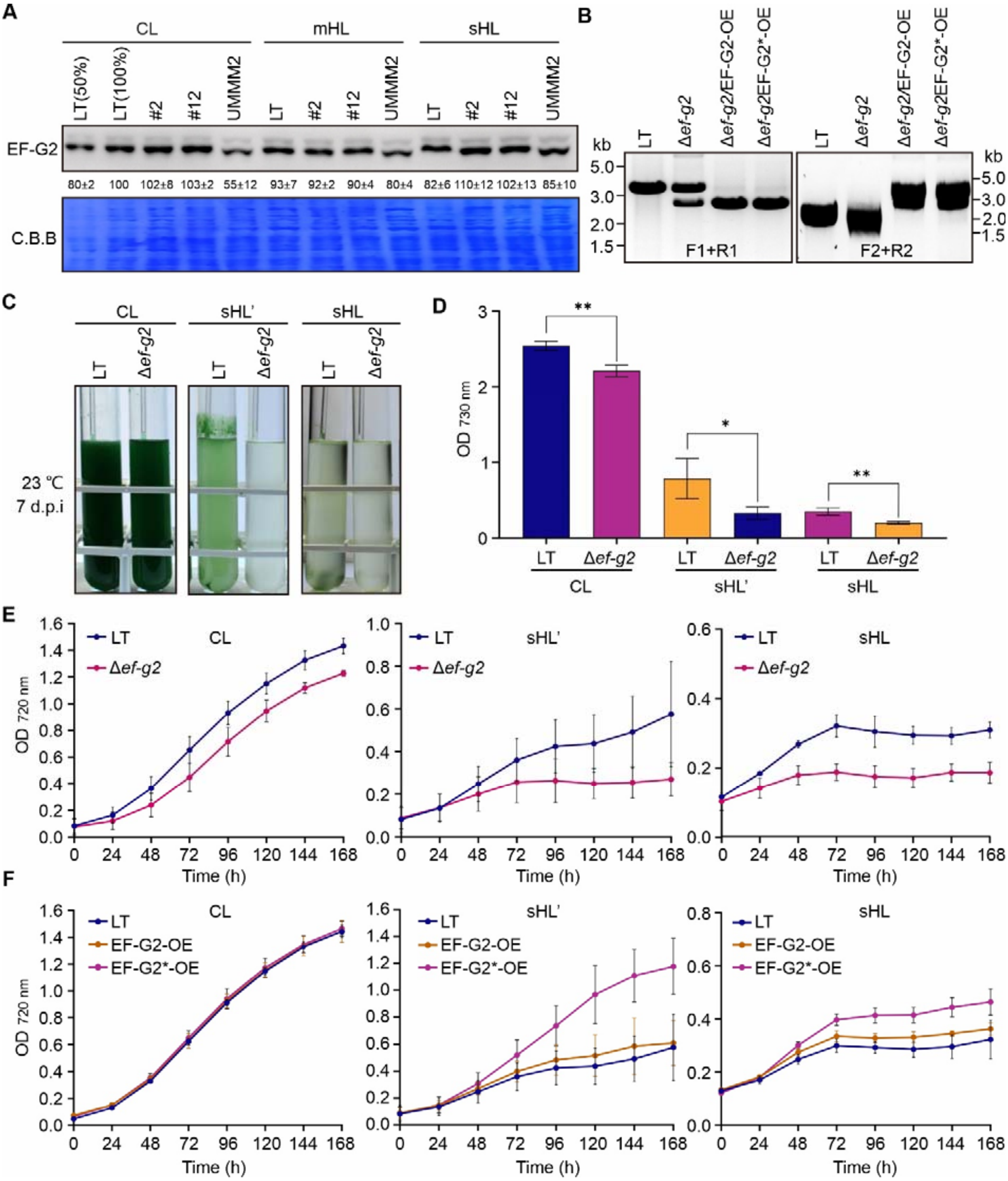


**Fig. S3.**

**Generation of *ef-g2/fusB* knockout and overexpression strains and growth analysis. (A)** Immunoblot analysis of EF-G2 in LT, EF-G2$_{R461C}$ (#2), NdhF1$_{F124L}$+EF-G2$_{R461C}$ (#12), and UMMM2 cells under three different light intensities. Coomassie Brilliant Blue (C.B.B.) staining of membranes served as a loading control. A representative result from 3 independent experiments is presented, along with average signal quantifications relative to LT (100%) ± SD. **(B)** PCR confirmation of complete segregation for KO and OE strains is displayed. *Δef-g2* represents KO of *ef-g2*/*fusB*, EF-G2-OE denotes OE of *ef-g2*, and EF-G2*-OE indicates OE of *ef-g2* with R461C mutation. **(C)** A representative image of LT and *Δef-g2* cultures grown for 7 days under different light intensities is shown. sHL' represents 1,000 µmol photons $m^{-2}$ $s^{-1}$. Pre-cultures were cultivated for 7 days at CL at 23°C. Three independent experiments were conducted. **(D)** Optical density ($OD_{730\ nm}$) of LT and *Δef-g2* on day 7 of cultivation is presented. Data represents mean ± SD from three independent experiments as in **C**. Statistical significance was determined using two-tailed Student's t-test, with * indicating $p < 0.05$ and ** indicating $p < 0.01$. **(E)** Growth curves of LT and *Δef-g2* under CL, sHL' and sHL conditions are shown. Data represents mean ± SD from three independent experiments as in **C**. **(F)** Growth curves of LT, EF-G2-OE, and EF-G2*-OE under CL, sHL' and sHL conditions are presented. Data represents mean ± SD from three independent experiments as in **Fig. 2F**.

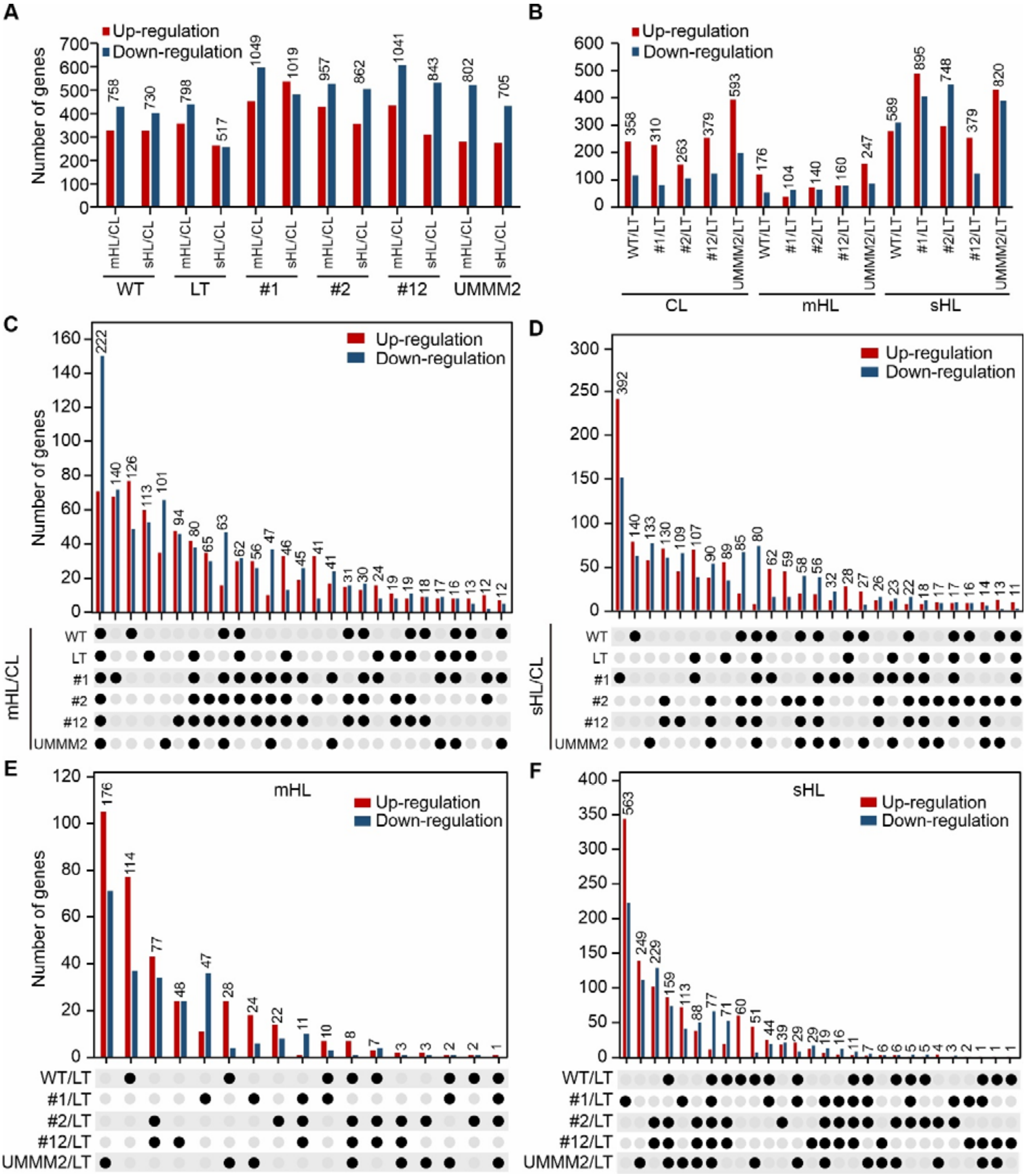


**Fig. S4.**

**Detailed transcriptomic analyses. (A)** This panel shows the number of DEGs for mHL vs. CL and sHL vs. CL in WT, LT and the four mutant strains: NdhF1$_{F124L}$ (#1), EF-G2$_{R461C}$ (#2), NdhF1$_{F124}$+EF-G2$_{R461C}$ (#12) and UMMM2. **(B)** The number of DEGs for WT and the four mutant strains vs. LT in CL, mHL and sHL conditions is presented. **(C-F)** Intersection (UpSet) plots show the number of commonly up- or down-regulated DEGs for various comparisons:

mHL vs. CL (**C**), sHL vs. CL (**D**) in different intersections of WT, LT and the four mutant strains, and for WT and the four mutant strains vs. LT under mHL (**E**) and sHL (**F**) conditions.

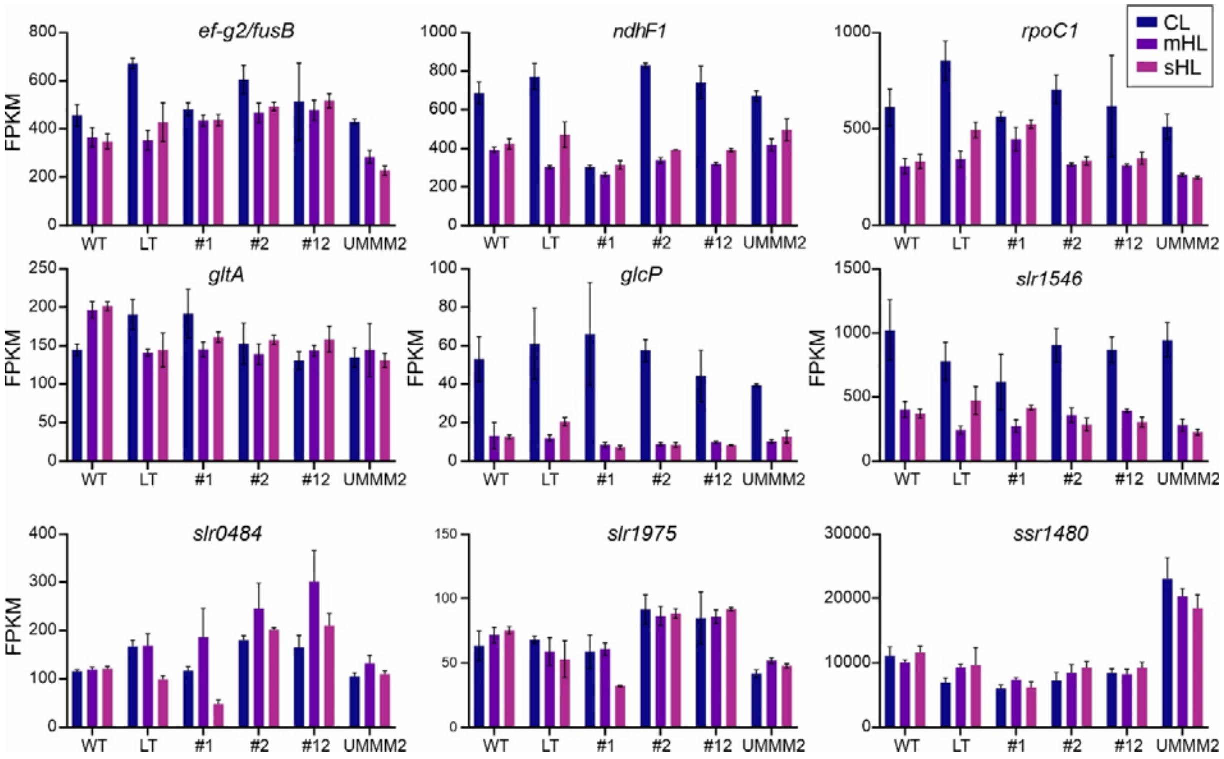


**Fig. S5.**

**Transcript abundance of genes mutated in UMMM2 in different genotypes and light conditions.** This figure displays transcript abundance (FPKM) for nine genes with high-frequency mutations in UMMM2, shown for WT, LT, NdhF1$_{F124L}$ (#1), EF-G2$_{R461C}$ (#2), NdhF1$_{F124L}$+EF-G2$_{R461C}$ (#12) and UMMM2 cells under three different light intensities.

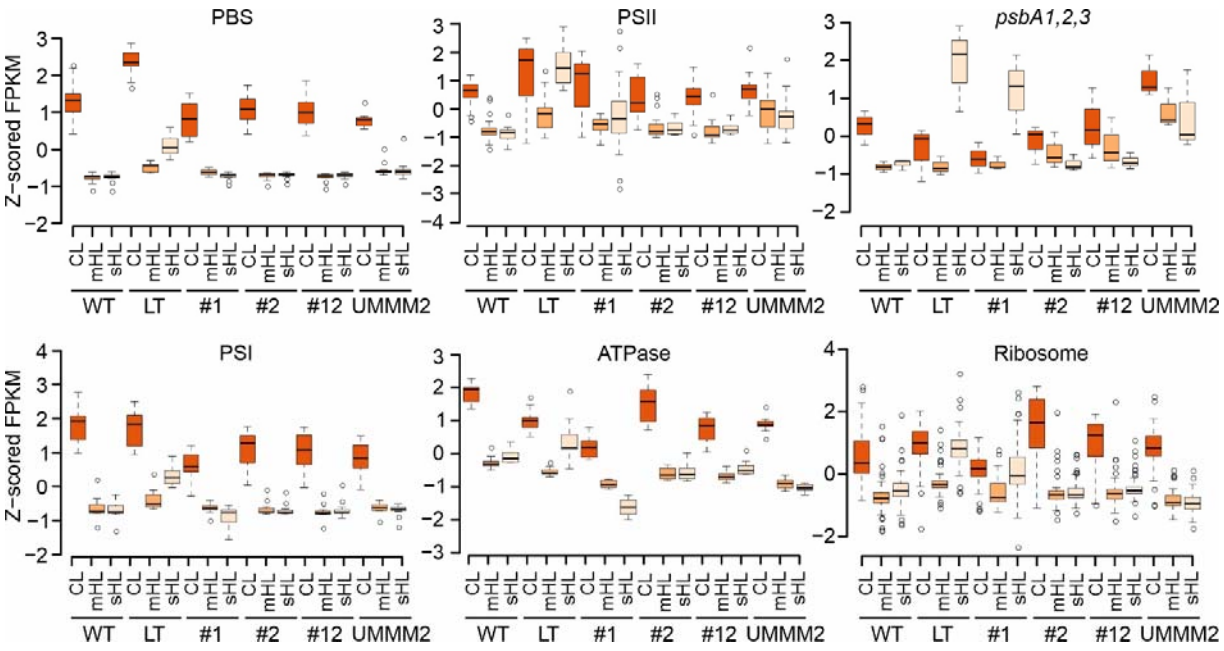


**Fig. S6.**

**Overview of transcriptional responses of photosynthesis- and ribosome-related genes in different genotypes and light conditions.** Z-scored FPKM values are presented for WT, LT, $NdhF1_{F124L}$ (#1), EF-$G2_{R461C}$ (#2), $NdhF1_{F124L}$+EF-$G2_{R461C}$ (#12) and UMMM2 cells under three different light intensities. The data covers five gene sets (all phycobilisome, PSII, PSI, ATP synthase, and ribosome subunits) and the three *psbA* genes.

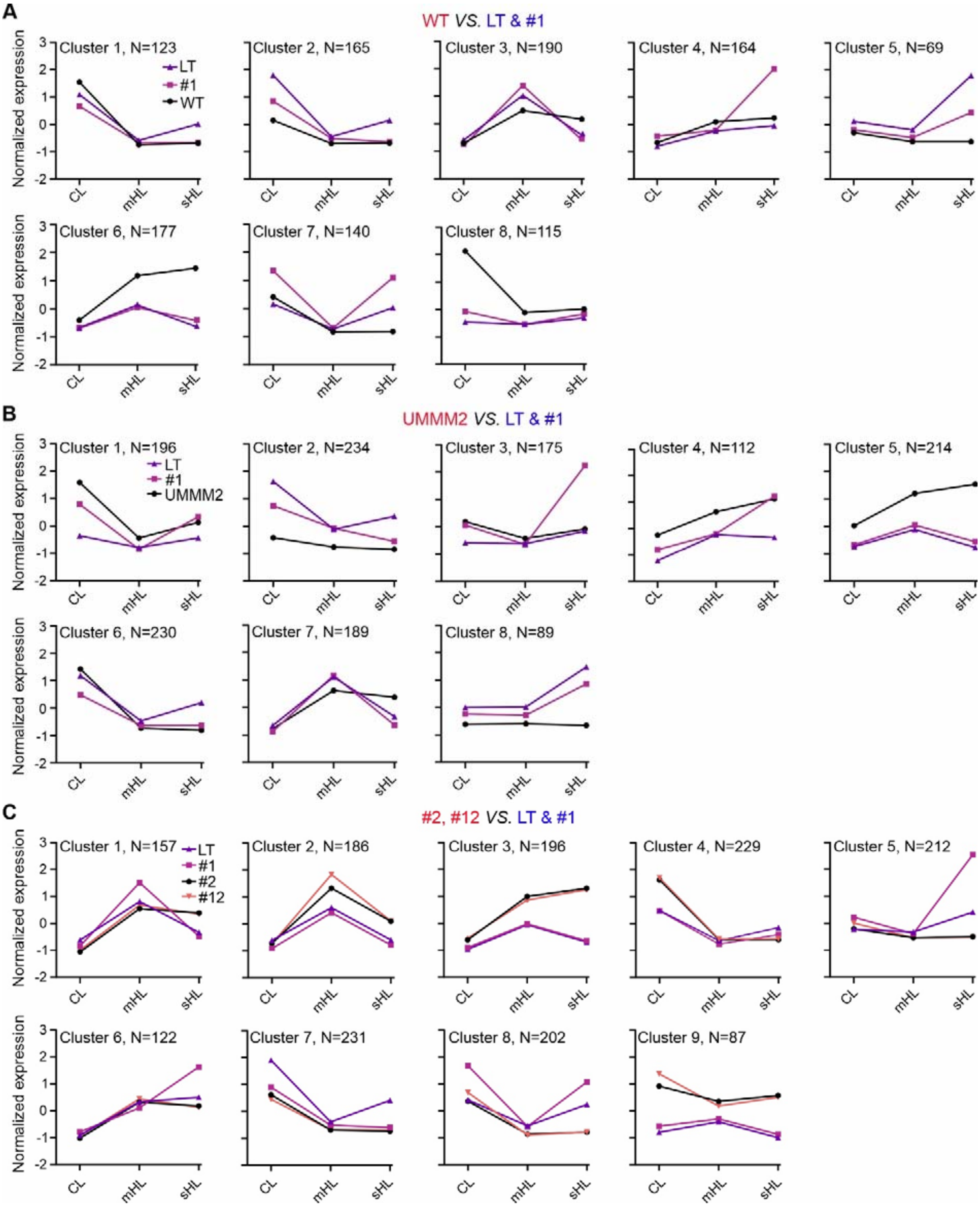


**Fig. S7.**

**Comparison of DEG clusters between HL-tolerant and -intolerant strains. (A-C)** Line plots show averaged gene expression patterns from Fuzzy c-means soft clustering analysis, comparing sHL-tolerant strains (WT, UMMM2, EF-G2$_{R461C}$ (#2), NdhF1$_{F124L}$+EF-G2$_{R461C}$ (#12)) with intolerant strains (LT, NdhF1$_{F124L}$ (#1)). Comparisons are shown for WT (**A**), UMMM2 (**B**), and #2 and #12 (**C**) against LT and #1. Gene numbers for each group are included. The x-axis represents light intensities (CL, mHL, sHL), and the y-axis shows mean normalized expression.

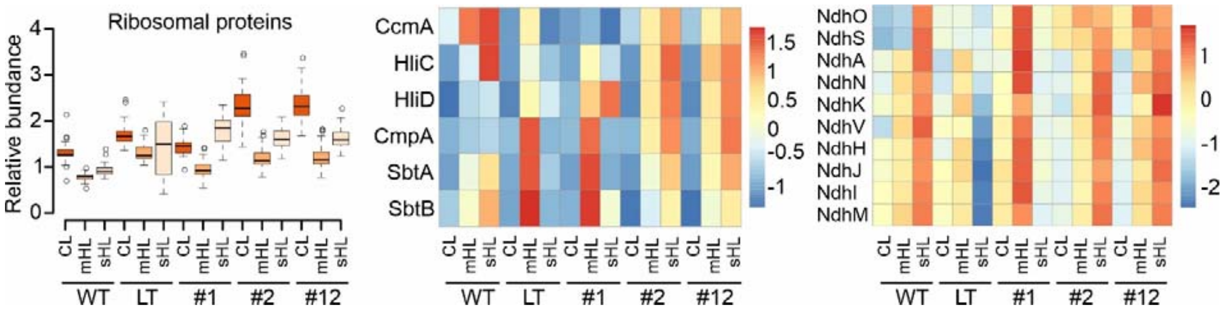


**Fig. S8.**

**Overview of specific protein set responses in different genotypes and light conditions.** Relative abundance values for WT, LT, UMMM2, NdhF1$_{F124L}$ (#1), EF-G2$_{R461C}$ (#2), and NdhF1$_{F124L}$+EF-G2$_{R461C}$ (#12) cells under three light intensities are shown for three protein sets: ribosome (Rpl1, 5-6, 9-13, 16-21, 23-24, 27, and Rps1, 5-7, 9-10, 12-14, 16, 19-20), Hlip/CCM, and NDH-1 proteins from regulon 12 in **Fig. 5**.

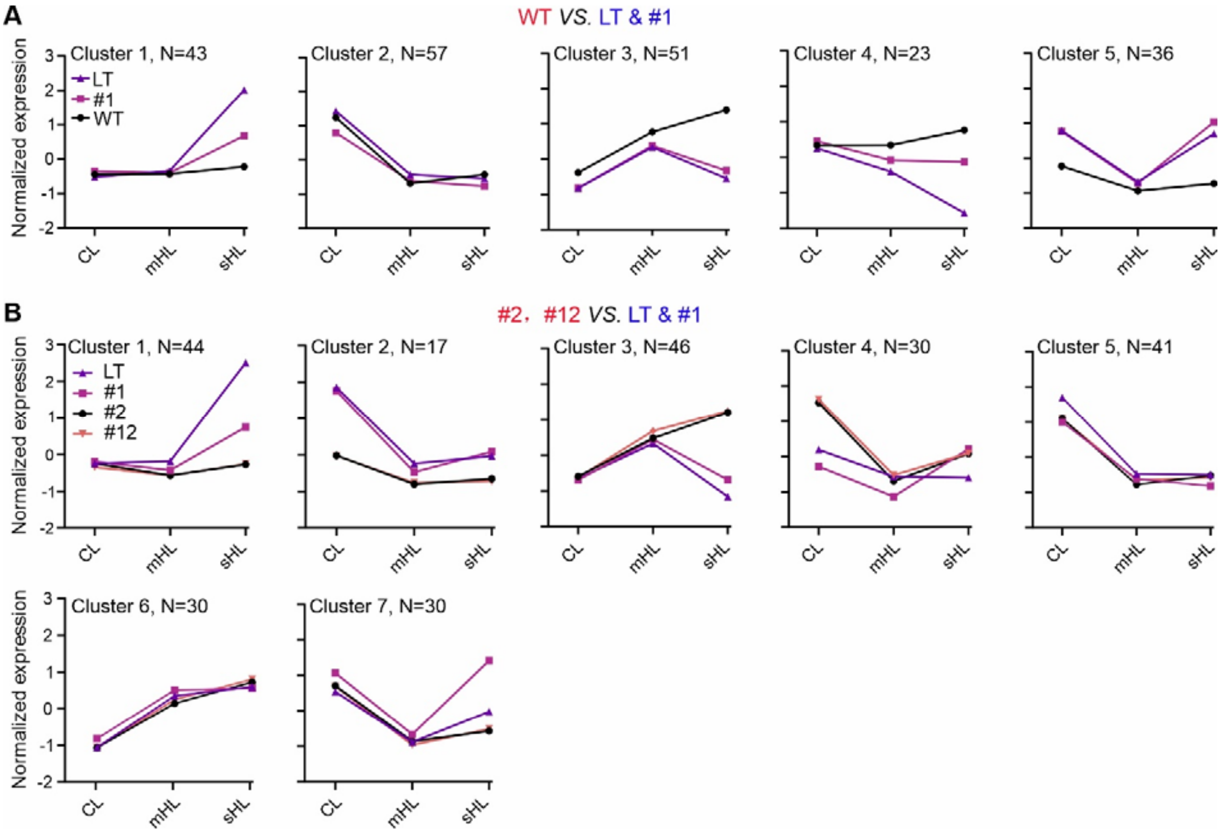


**Fig. S9.**

**Comparison of DEP clusters between HL-tolerant and -intolerant strains. (A, B)** Line plots display averaged protein expression patterns from Fuzzy c-means soft clustering analysis, comparing sHL-tolerant strains (WT, EF-G2$_{R461C}$ (#2), NdhF1$_{F124L}$+EF-G2$_{R461C}$ (#12)) with intolerant strains (LT, Ndh$_{F1F124L}$ (#1)). Comparisons for WT (**A**) and #2 and #12 (**B**) against LT and #1 are shown. Protein numbers for each group are included. The x-axis represents light intensities (CL, mHL, sHL), and the y-axis shows mean normalized expression.

**A**

| Log2 | mHL | sHL |
|---|---|---|
| WT/LT | $p$ = 1.23e-56, $\hat{r}_{Pearson}$ = 0.56 | $p$ = 5.13e-19, $\hat{r}_{Pearson}$ = 0.34 |
| #1/LT | $p$ = 1.57e-36, $\hat{r}_{Pearson}$ = 0.46 | $p$ = 6.30e-11, $\hat{r}_{Pearson}$ = 0.25 |
| #2/LT | $p$ = 2.01e-19, $\hat{r}_{Pearson}$ = 0.34 | $p$ = 2.56e-14, $\hat{r}_{Pearson}$ = 0.29 |
| #12/LT | $p$ = 9.08e-15, $\hat{r}_{Pearson}$ = 0.30 | $p$ = 2.87e-15, $\hat{r}_{Pearson}$ = 0.30 |

**B**

| | Log2(mHL/CL) | Log2(sHL/CL) |
|---|---|---|
| WT | $p$ = 4.57e-86, $\hat{r}_{Pearson}$ = 0.66 | $p$ = 3.58e-77, $\hat{r}_{Pearson}$ = 0.64 |
| LT | $p$ = 8.35e-53, $\hat{r}_{Pearson}$ = 0.54 | $p$ = 1.08e-07, $\hat{r}_{Pearson}$ = 0.20 |
| #1 | $p$ = 7.09e-56, $\hat{r}_{Pearson}$ = 0.56 | $p$ = 1.14e-39, $\hat{r}_{Pearson}$ = 0.48 |
| #2 | $p$ = 6.04e-87, $\hat{r}_{Pearson}$ = 0.67 | $p$ = 4.86e-83, $\hat{r}_{Pearson}$ = 0.66 |
| #12 | $p$ = 6.04e-87, $\hat{r}_{Pearson}$ = 0.67 | $p$ = 5.396e-90, $\hat{r}_{Pearson}$ = 0.68 |

**Fig. S10.**

**Comparison of transcriptome and proteome responses. (A, B)** Correlation analysis between transcriptomes and proteomes. Pearson correlation coefficients (r) and Student t-test p-values (p) are provided for expression changes between all transcript-protein pairs of WT, NdhF1$_{F124L}$ (#1), EF-G2$_{R461C}$ (#2), and NdhF1$_{F124L}$+EF-G2$_{R461C}$ (#12) relative to LT under mHL and sHL conditions (**A**), and between all transcript-protein pairs of mHL and sHL relative to CL for the five genotypes (**B**).

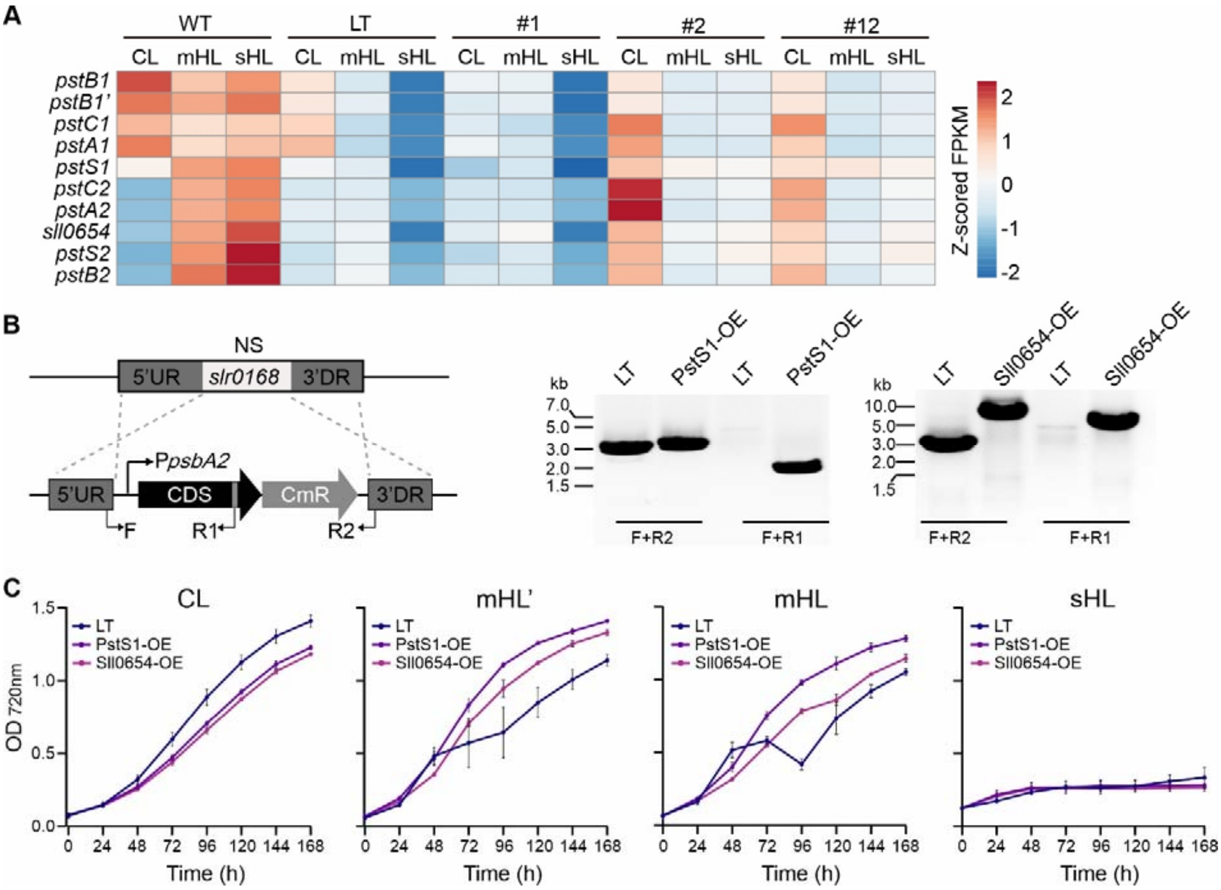


**Fig. S11.**

**Generation of overexpression strains and growth analysis.** **(A)** Heatmap of Z-score normalized FPKM for Pho regulon transcripts, comparing sHL-tolerant strains (WT, EF-G2$_{R461C}$ (#2), NdhF1$_{F124L}$+EF-G2$_{R461C}$ (#12)) with sHL-intolerant strains (LT, NdhF1$_{F124L}$ (#1)) under three light conditions. **(B)** PCR confirmation of complete segregation in overexpression strains, using the strategy illustrated in **fig. S1A**. **(C)** Growth curves of LT, PstS1-OE, and Sll0654-OE under CL, mHL', mHL, and sHL conditions (mHL': 500 µmol photons $m^{-2}$ $s^{-1}$). Data shows mean ± SD from three independent experiments as in **Fig. 9**.

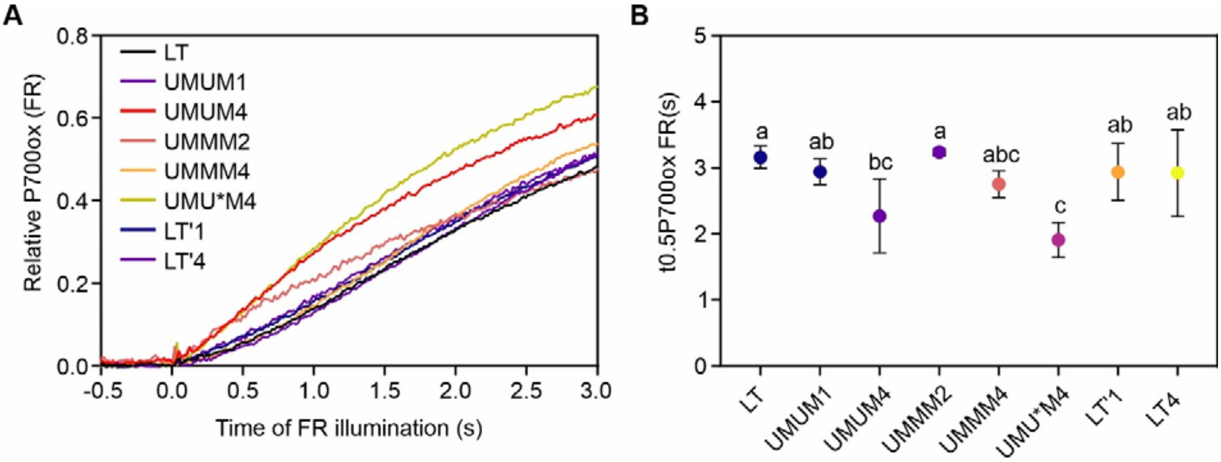


**Fig. S12.**

**Measurement of CEF in HL-evolved strains. (A)** P700 oxidation kinetics of LT, #1, and HL-evolved strains. **(B)** Half-times of P700 oxidation, showing mean ± SD from four independent experiments as in **Fig. 10B**. Different letters above error bars indicate statistical differences ($p < 0.05$) determined by one-way ANOVA with post-hoc Tukey HSD test.

**Supplementary Table 1** Common HL-tolerance related genes (HL-DEGs) in the HL-tolerant strains. Commonly identified up- and down-regulated HL-DEGs in HL-tolerant WT, UMMM2, and EF-G2$_{R461C}$ /NdhF1$_{F124L}$+EF-G2$_{R461C}$ as showed in **Figure 4D** are listed.

| Gene ID | Old locus tag | Name | Description |
|---|---|---|---|
| | | **Down_regulated HL-DEGs** | |
| SGL_RS16410 | sll0743 | sll0743 | hypothetical protein |
| SGL_RS16215 | slr0699 | slr0699 | unknown protein |
| SGL_RS08830 | sll1596 | kaiB2 | circadian clock protein KaiB homolog |
| SGL_RS08295 | slr0870 | slr0870 | hypothetical protein |
| SGL_RS05160 | slr1396 | slr1396 | unknown protein |
| SGL_RS01830 | slr6101 | slr6101 | plasmid-encoded putative toxin/antitoxin pair |
| SGL_RS01700 | slr6072 | slr6072 | unknown protein |
| SGL_RS01575 | slr6045 | slr6045 | unknown protein |
| SGL_RS01430 | slr6013 | slr6013 | unknown protein |
| SGL_RS01030 | sll7086 | sll7086 | unknown protein |
| SGL_RS01025 | sll7085 | sll7085 | unknown protein |
| SGL_RS00605 | sll5128 | sll5128 | unknown protein |
| SGL_RS09525 | | | hypothetical protein |
| SGL_RS14650 | | | type II toxin-antitoxin system HicA family |
| | | **Up_regulated HL-DEGs** | |
| SGL_RS16520 | sll0037 | cbiX | sirohydrochlorin cobaltochelatase |
| SGL_RS12205 | sll0141 | sll0141 | putative periplasmic adaptor protein |
| SGL_RS13285 | sll0314 | sll0314 | periplasmic protein, function unknown |
| SGL_RS14445 | sll0381 | sll0381 | hypothetical protein |
| SGL_RS14430 | sll0384 | sll0384 | cation and iron carrying protein |
| SGL_RS14425 | sll0385 | sll0385 | ATP-binding protein of ABC transporter |
| SGL_RS15490 | sll0477 | sll0477 | putative biopolymer transport ExbB-like protein |
| SGL_RS14310 | sll0608 | ycf49 | hypothetical protein YCF49 |
| SGL_RS03825 | sll0656 | nucH | extracellular nuclease |
| SGL_RS17785 | sll0720 | sll0720 | RTX toxin activating protein homolog |
| SGL_RS17775 | sll0722 | sll0722 | unknown protein |
| SGL_RS17765 | sll0723 | sll0723 | unknown protein |
| SGL_RS07350 | sll1308 | sll1308 | probable oxidoreductase |
| SGL_RS17545 | sll1483 | sll1483 | periplasmic protein |
| SGL_RS04105 | sll1507 | sll1507 | salt-induced periplasmic protein |
| SGL_RS17790 | sll1552 | sll1552 | unknown protein |
| SGL_RS06130 | sll1968 | pmgA | photomixotrophic growth related protein |
| SGL_RS00225 | sll5050 | sll5050 | probable glycosyltransferase |
| SGL_RS00580 | sll5123 | sll5123 | SOS mutagenesis and repair, UmuD protein homolog |
| SGL_RS01415 | sll6010 | sll6010 | unknown protein |
| SGL_RS01685 | sll6069 | sll6069 | unknown protein |
| SGL_RS14105 | slr0492 | menE | O-succinylbenzoic acid-CoA ligase |
| SGL_RS15495 | slr0513 | futA2 | iron transport system substrate-binding protein |
| SGL_RS15505 | slr0516 | slr0516 | hypothetical protein |
| SGL_RS02010 | slr1485 | slr1485 | putative phosphatidylinositol phosphate kinase |
| SGL_RS09605 | slr2131 | acrB | RND multidrug efflux transporter |
| SGL_RS00245 | slr5054 | slr5054 | probable glycosyltransferase |
| SGL_RS01405 | slr6008 | slr6008 | unknown protein |
| SGL_RS00880 | ssl7053 | ssl7053 | hypothetical protein |
| SGL_RS04605 | ssr3465 | ssr3465 | unknown protein |
| SGL_RS19830 | | | hypothetical protein |

**Supplementary Table 2** Common HL-tolerance related proteins (HL-DEPs) in the HL-tolerant strains. Commonly identified up- and down-regulated HL-DEPs in HL-tolerant WT, and EF-G2$_{R461C}$ /NdhF1$_{F124L}$+EF-G2$_{R461C}$ as showed in **Figure 6C** are listed.

| Protein ID | Name | Uniprot ID | Description |
|---|---|---|---|
| | | **Down_regulated HL-DEPs** | |
| Sll0172 | Sll0172 | Q55558 | Periplasmic protein, function unknown |
| Sll0258 | PsbV | Q55013 | Cytochrome c550 |
| Sll0293 | Sll0293 | Q55547 | Unknown protein |
| Sll0915 | PqqE | P74305 | Periplasmic protease |
| Sll1491 | Sll1491 | P74598 | Periplasmic WD-repeat protein |
| Sll1835 | Sll1835 | P73111 | Periplasmic protein, function unknown |
| Slr1266 | Slr1266 | P74179 | Hypothetical protein |
| Slr1274 | PilM | P74186 | Probable fimbrial assembly protein PilM |
| Slr1276 | Slr1276 | P74188 | Hypothetical protein |
| | | **Up_regulated HL-DEPs** | |
| Sll1394 | MsrA1 | P72622 | Peptide methionine sulfoxide reductase |
| Slr1367 | GlgP | P73546 | Glycogen phosphorylase |
| Sll1961 | Sll1961 | P73804 | Hypothetical protein |
| Sll0553 | Sll0553 | Q55390 | Hypothetical protein |
| Slr0453 | Slr0453 | P74690 | Hypothetical protein |
| Slr2075 | GroES | Q05971 | 10 kD chaperonin |
| Sll1294 | PilJ | P73173 | Methyl-accepting chemotaxis protein |
| slr0607 | Slr0607 | P74754 | Hypothetical protein |
| ssl1633 | HliC, scpB | P73563 | High light-inducible polypeptide HliC |
| Slr0884 | Gap1 | P49433 | Glyceraldehyde 3-phosphate dehydrogenase 1 |
| Sll0680 | PstS1 | Q55199 | Periplasmic phosphate-binding protein |
| Sll1594 | NdhR | P73862 | NdhF3 operon transcriptional regulator |
| Sll0934 | CcmA | P72864 | Carboxysome formation protein CcmA |
| Sll0329 | Gnd | P52208 | 6-phosphogluconate dehydrogenase |
| Sll0654 | Sll0654 | P72939 | Alkaline phosphatase |
| Slr0516 | Slr0516 | Q55837 | Hypothetical protein |
| Slr1351 | MurF | P45450 | UDP-N-acetylmuramoyl-pentapeptide synthase |
| Slr1512 | SbtA | P73953 | Sodium-dependent bicarbonate transporter |
| Slr1204 | HtrA | P73354 | Protease |
| Slr2073 | Ycf50 | P73376 | Hypothetical protein YCF50 |
| Slr1280 | NdhK | P19050 | NADH dehydrogenase subunit NdhK |
| Sll1680 | Sll1680 | P72779 | Hypothetical protein |
| Sll1566 | GgpS | P74258 | Glucosylglycerolphosphate synthase |
| Sll0185 | Sll0185 | Q55770 | Hypothetical protein |
| Sll0381 | Sll0381 | Q55744 | Hypothetical protein |
